# Genome-wide association analyses of individual differences in quantitatively assessed reading- and language-related skills in up to 34,000 people

**DOI:** 10.1101/2021.11.04.466897

**Authors:** Else Eising, Nazanin Mirza-Schreiber, Eveline L. de Zeeuw, Carol A. Wang, Dongnhu T. Truong, Andrea G. Allegrini, Chin Yang Shapland, Gu Zhu, Karen G. Wigg, Margot Gerritse, Barbara Molz, Gökberk Alagöz, Alessandro Gialluisi, Filippo Abbondanza, Kaili Rimfeld, Marjolein van Donkelaar, Zhijie Liao, Philip R. Jansen, Till F. M. Andlauer, Timothy C. Bates, Manon Bernard, Kirsten Blokland, Anders D. Børglum, Thomas Bourgeron, Daniel Brandeis, Fabiola Ceroni, Philip S. Dale, Karin Landerl, Heikki Lyytinen, Peter F. de Jong, John C. DeFries, Ditte Demontis, Yu Feng, Scott D. Gordon, Sharon L. Guger, Marianna E. Hayiou-Thomas, Juan A. Hernández-Cabrera, Jouke-Jan Hottenga, Charles Hulme, Elizabeth N. Kerr, Tanner Koomar, Maureen W. Lovett, Nicholas G. Martin, Angela Martinelli, Urs Maurer, Jacob J. Michaelson, Kristina Moll, Anthony P. Monaco, Angela T. Morgan, Markus M. Nöthen, Zdenka Pausova, Craig E. Pennell, Bruce F Pennington, Kaitlyn M. Price, Veera M. Rajagopal, Frank Ramus, Louis Richer, Nuala H. Simpson, Shelley Smith, Margaret J. Snowling, John Stein, Lisa J. Strug, Joel B. Talcott, Henning Tiemeier, Marc M.P. van de Schroeff, Ellen Verhoef, Kate E. Watkins, Margaret Wilkinson, Margaret J. Wright, Cathy L. Barr, Dorret I. Boomsma, Manuel Carreiras, Marie-Christine J. Franken, Jeffrey R. Gruen, Michelle Luciano, Bertram Müller-Myhsok, Dianne F. Newbury, Richard K. Olson, Silvia Paracchini, Tomas Paus, Robert Plomin, Gerd Schulte-Körne, Sheena Reilly, J. Bruce Tomblin, Elsje van Bergen, Andrew J.O. Whitehouse, Erik G. Willcutt, Beate St Pourcain, Clyde Francks, Simon E. Fisher

## Abstract

The use of spoken and written language is a capacity that is unique to humans. Individual differences in reading- and language-related skills are influenced by genetic variation, with twin-based heritability estimates of 30-80%, depending on the trait. The relevant genetic architecture is complex, heterogeneous, and multifactorial, and yet to be investigated with well-powered studies. Here, we present a multicohort genome-wide association study (GWAS) of five traits assessed individually using psychometric measures: word reading, nonword reading, spelling, phoneme awareness, and nonword repetition, with total sample sizes ranging from 13,633 to 33,959 participants aged 5-26 years (12,411 to 27,180 for those with European ancestry, defined by principal component analyses). We identified a genome-wide significant association with word reading (rs11208009, p=1.098 × 10^−8^) independent of known loci associated with intelligence or educational attainment. All five reading-/language-related traits had robust SNP-heritability estimates (0.13–0.26), and genetic correlations between them were modest to high. Using genomic structural equation modelling, we found evidence for a shared genetic factor explaining the majority of variation in word and nonword reading, spelling, and phoneme awareness, which only partially overlapped with genetic variation contributing to nonword repetition, intelligence and educational attainment. A multivariate GWAS was performed to jointly analyse word and nonword reading, spelling, and phoneme awareness, maximizing power for follow-up investigation. Genetic correlation analysis of multivariate GWAS results with neuroimaging traits identified association with cortical surface area of the banks of the left superior temporal sulcus, a brain region with known links to processing of spoken and written language. Analysis of evolutionary annotations on the lineage that led to modern humans showed enriched heritability in regions depleted of Neanderthal variants. Together, these results provide new avenues for deciphering the biological underpinnings of these uniquely human traits.

## Introduction

The processing and production of complex spoken and written language are capacities that appear to be distinct to our species. Such skills have become fundamental for day-to-day life in modern society. Decades of family and twin studies have revealed substantial genetic components contributing to individual variation in reading- and language-related traits, as well as to susceptibility to associated disorders. A recent meta-analysis integrated available data on these skills from 49 twin studies, with a total sample size of 38,000 children and adolescents, aged 4-18 years. The meta-analysis yielded heritability estimates of 66% for word reading (meta-analysis of 48 studies), 80% for spelling (15 studies), and 52% for phoneme awareness (the ability to identify and manipulate individual sounds of spoken words; 13 studies), and suggested greater genetic influences on reading-related abilities than language-related measures (heritability 34%; meta-analysis of 10 studies with measures on receptive and expressive vocabulary, oral language and naming abilities)^1^.

Linkage mapping and candidate gene studies have reported associations of single-nucleotide polymorphisms (SNPs) and/or genetic loci with reading and language-related traits, as well as with disorders such as dyslexia and developmental language disorder (DLD; which encompasses the older definition of specific language impairment or SLI)^2^. However, replication efforts have met with limited success. Genome-wide association studies (GWAS) are beginning to identify SNPs that show genome-wide significant associations with reading- and language-related traits: rs7642482 near *ROBO2* associated with expressive vocabulary in infancy^3^, rs17663182 within MIR924HG with rapid automatized naming of letters^4^, and rs1555839 near *RPL7P34* with rapid automatized naming and rapid alternating stimulus, deficits of which are often implicated in dyslexia^5^. Nonetheless, insights into the genomic underpinnings of these types of skills from GWAS approaches have thus far been limited, which may reflect low power due to the relatively small sample sizes of the cohorts, such that the majority of genetic variance remains unexplained. Sample sizes have remained limited because of the labour-intensive assessment methods required for phenotyping reading- and language-related traits, which are difficult or even impossible to replace with simple questionnaires. Yet, well-powered GWAS efforts that characterize the molecular genetic variation involved in reading- and language-related traits can provide novel perspectives on the biological bases and origins of human cognitive specializations^6^.

Here, we present large-scale GWAS meta-analyses of a set of reading- and language-related traits, measured with psychometric tools. We captured variation across the phenotypic spectrum, extending beyond disorder. Our study focused on traits that have been assessed using continuous measures in multiple cohorts from the international GenLang network (www.genlang.org), together with several public datasets that have data available for the relevant phenotypes, matched to genome-wide genotype information. Five quantitative traits were identified for which phenotype data could be aligned across different cohorts, to yield sufficiently large sample sizes for GWAS: word reading, nonword reading, spelling, phoneme awareness and nonword repetition. Univariate GWAS meta-analyses were performed to identify genetic variation influencing these traits and to model the genetic overlaps between them. For comparative purposes, a GWAS meta-analysis for performance IQ was also performed in the same dataset. Together with publicly available GWAS summary statistics from prior studies of cognitive performance and educational attainment, these data were used to study genetic relationships between reading- and language-related traits, IQ and educational attainment. A multivariate approach allowed us to optimise the power of the word reading GWAS meta-analysis for functional follow-ups investigating the tissues, cell types, brain regions and evolutionary signatures involved.

## Results

### GWAS meta-analysis for quantitative reading- and language-related traits

We studied five quantitative reading- and language-related traits: word reading accuracy, nonword reading accuracy, spelling accuracy, phoneme awareness and nonword repetition accuracy (Table 1). These traits are thought to tap into a number of underlying processes involved in written and spoken language. For example, nonword reading relies heavily on basic decoding skills: translating graphemes one by one into phonemes^7^, while spelling utilizes lexical and orthographic knowledge: understanding of permissible letter patterns and how they are arranged in specific words^8^. Phoneme awareness measures the ability to distinguish and manipulate the separate phonemes in spoken words^9^. Nonword repetition tasks tap into speech perception, phonological short term memory and articulation^10^. A total of 22 cohorts, aggregated by the GenLang consortium combined with several publicly available datasets, provided data for one or several of these traits (Supplemental Table 1 and 2, Supplemental Figure 1 and 2). The cohorts connected in the GenLang network have either been originally ascertained through a proband with language/reading disorder (DLD/SLI or dyslexia) or were sampled from the general population; all cohorts include quantitative phenotypic data gathered via validated psychometric tests. Some of the samples are birth cohorts, and some involve family or twin designs. The phenotype data were collected across an array of different ages, test instruments and languages (primarily English, and also Dutch, Spanish, German, French, Finnish, and Hungarian). We reduced heterogeneity of assessment age by excluding individuals over 18 years of age (except for three cohorts, see Supplemental Methods), and, where phenotype data were available from the same participant at multiple ages, by choosing the age that matched best with the assessment ages of the largest cohort(s). We limited the heterogeneity introduced by different test instruments by only including those that measured the phenotypes described in our analysis plan (Supplemental Notes), and, in cases where data from more than one test instrument were available, by selecting the test instrument that was used by the largest number of cohorts.

**Table 1:** Phenotypes and sample sizes of the GWAS meta-analyses.

| Trait | Phenotype description | Meta-analysis total sample |  | Meta-analysis European ancestry only |  |
| --- | --- | --- | --- | --- | --- |
|  |  | # cohorts | # individuals | # cohorts | # individuals |
| Word reading | Number of correct words read aloud from a list in a time restricted or unrestricted fashion | 19 | 33,959 | 18 | 27,180 |
| Nonword* reading | Number of nonwords read aloud correctly from a list in a time restricted or unrestricted fashion | 13 | 17,984 | 12 | 16,746 |
| Spelling | Number of words correctly spelled orally or in writing, after being dictated as single words or in a sentence | 15 | 18,514 | 14 | 17,278 |
| Phoneme awareness | Number of words correctly altered in phoneme deletion/elision and spoonerism tasks | 12 | 13,633 | 11 | 12,411 |
| Nonword* repetition | Number of nonwords or phonemes repeated aloud correctly | 10 | 14,046 | 10 | 12,828 |

\*A nonword is a group of phonemes that looks or sounds like a word, obeys the phonotactic rules of the language, but has no meaning.

GWAS meta-analysis results for word reading and nonword reading stratified by age or test instrument were used to investigate the remaining amount of heterogeneity related to age and use of different test instruments (Supplemental Table 3). Genetic correlations calculated with linkage disequilibrium score regression (LDSC)^11^ between age-stratified meta-analysis results (cohorts with mean age <12 years vs ≥12 years; see Supplemental Table 1) were high (word reading rg=0.86, se=0.16; nonword reading rg=0.88, se=0.24). GWAS meta-analysis results of the most commonly used reading test (Time Limited Word Recognition Test; TOWRE) were found to be highly correlated with GWAS meta-analysis results of all other reading tests (rg=0.85, se=0.15 for word reading, rg=0.99, se=0.21 for nonword reading). Similar results were obtained when comparing GWAS meta-analysis results of time-restricted reading tests to all other reading tests. Limited heterogeneity of GWAS meta-analysis results was also evident in the Cochran Q statistics (Supplemental Figure 3) and LDSC ratios (Supplemental Table 4) for all traits except nonword repetition.

We also assessed whether there was heterogeneity related to sex effects, by estimating genetic correlations for female- and male-only subsets. Results of male and female subsets were highly correlated (estimates of genetic correlation are rg=1.04, se=0.17, p=1.7×10^−9^ for word reading, rg=0.97, se=0.19, p=2.1×10^−7^ for nonword reading, rg=1.11, se=0.29, p=1.0×10^−4^ for spelling, rg=1.56, se=0.55, p=4.2×10^−3^ for phoneme awareness and rg=1.30, se=0.63, p=0.039 for nonword repetition; some estimates are above 1, because the genetic covariance estimator is not constrained in LDSC, and likely represents sampling variation and randomness; Supplemental Table 3).

### A genome-wide significant locus associated with word reading

For evaluating statistical significance of SNP associations, we determined an appropriate threshold that was adjusted for multiple-testing based on the correlation structure of our five reading-/language-related traits, as estimated using phenoSpD. The significance threshold for assessing the GWAS meta-analysis results was thus set to 5×10^−8^ / 2.15 independent traits = 2.33×10^−8^. We identified a genome-wide significant locus associated with word reading (rs11208009 C/T on chromosome 1, p=1.10×10^−8^) (Supplemental Figure 5). Rs11208009 has not shown association with general cognitive performance (IQ) or educational attainment, but other SNPs in LD with rs11208009 (r^2^>0.6) have been associated with triglyceride and total cholesterol levels in blood in previous GWAS studies (Supplemental Table 5). Three genes are located in the vicinity of rs11208009 and SNPs in LD (r^2^>0.6): *DOCK7*, encoding a guanine nucleotide exchange factor important for neurogenesis^12^; *ANGPTL3*, which encodes a growth factor specific for the vascular endothelium that is expressed specifically in the liver^13^; and *USP1*, encoding a deubiquitinating enzyme specific for the Fanconi anemia pathway^14^. The associated locus harbours an eQTL regulating *DOCK7* and *ATG4C* (another nearby gene which encodes an autophagy regulator^15^) in the cerebellum, and *DOCK7, ATG4C* and *USP1* in blood samples (Supplemental Table 6). Genome-wide significant loci were not identified for the other traits. Supplemental Table 7 lists all results with p<1×10^−6^.

### Traits related to reading and language are highly correlated at the genetic level

All five traits showed significant SNP heritability, with LDSC-based estimates ranging from 0.13 for nonword repetition to 0.26 for nonword reading (Supplemental Table 4), indicating that the captured common genetic variation accounts for a substantive proportion of the phenotypic variance in these skills. Pairwise genetic correlation analyses showed significant overlap among the reading- and language-related traits (Figure 1A; Supplemental Table 4). Genetic correlation estimates were especially high for word reading, nonword reading, spelling and phoneme awareness, ranging from 0.96 (se=0.07) to 1.06 (se=0.07).

**Figure 1:**
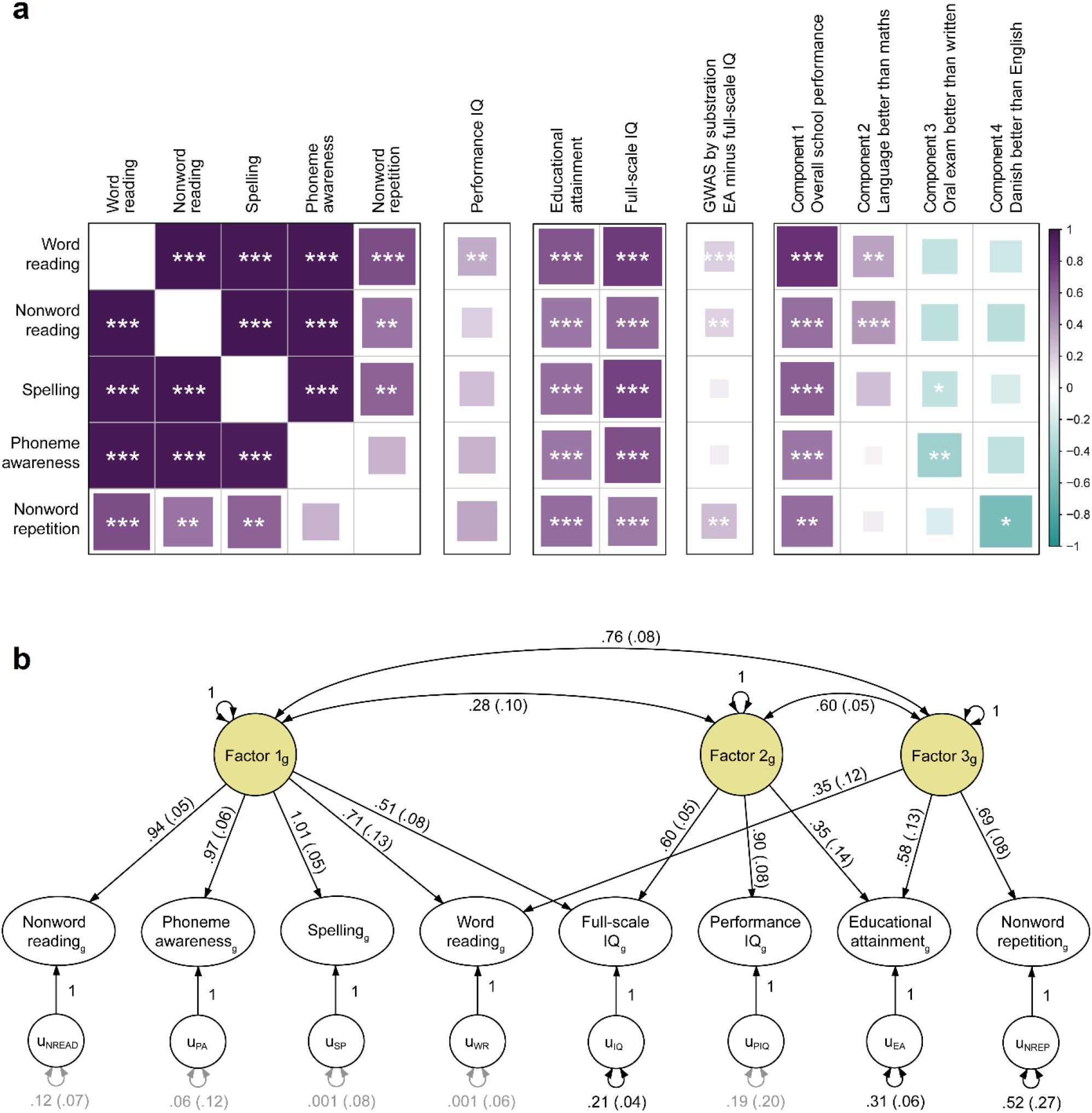
Reading- and language-related traits have a shared genetic architecture that is largely independent of performance IQ. **A**: Genetic correlations (rg) among the reading- and language-related traits, estimated with LDSC. Estimates capped at 1. Full LDSC results are reported in Supplemental Table 4. In addition, genetic correlations are given between the GenLang traits and 1) performance IQ (using GenLang cohorts only), 2) educational attainment (n=766,345) and full-scale IQ (n=257,828)^16^, 3) non-cognitive abilities involved in educational attainment, resulting from a recent GWAS-by-subtraction study (n=510,795)^17^, and 4) components associated with distinct performance domains, identified used a decomposition analysis of Danish school grades (n=30,982)^18^. Full results can be found in Supplemental Table 8. *Significant genetic correlation after correction for 18.28 independent comparisons (p<2.74×10^−3^); **p<2.74×10^−4^; ***p<2.74×10^−5^. **B**: Three factor model fitted to the GenLang summary statistics for word reading, nonword reading, spelling, phoneme awareness and nonword repetition and performance IQ, and published GWAS summary statistics for full-scale IQ and educational attainment^16^ using GenomicSEM^19^. Black and grey paths represent factor loadings with p<0.05 and p>0.05, respectively. Standardized factor loadings are shown, with standard error in parentheses. The subscript g represents the genetic variables; the u variables represent the residual genetic variance not explained by the models. Unstandardized results and model fit indices are reported in Supplemental Table 9.

Extensive prior literature has shown correlations of reading- and language-related traits with general cognitive performance and educational attainment. Most cognitive assessments depend on a combination of verbal and nonverbal tests. To enable the investigation of genetic overlaps between non-verbal cognitive performance and reading- and language-related traits, while closely matching the sample characteristics of our study, we carried out a GWAS meta-analysis of performance IQ in the GenLang network (n=18,722; Supplemental Figure 1-3). Only non-verbal subtests of general intelligence tests were used in this analysis (Supplemental Table 1). Summary statistics were also obtained from another three sources: 1) genome-wide studies of full-scale IQ (n=257,828) and educational attainment (n=766,345) by the Social Science Genetic Association Consortium^16^; 2) a GWAS-by-subtraction study that investigated the non-cognitive abilities involved in educational attainment (n=510,795)^17^; and 3) a recent GWAS analysis of school grades in the Danish iPSYCH cohort (n=30,982), that used a decomposition analysis to identify genetic associations with distinct domains of performance^18^.

The five GenLang traits showed moderate to strong positive genetic correlations with full-scale IQ (range 0.52–0.77), educational attainment (range 0.54–0.68) and school performance (range 0.54– 0.81) (Figure 1A, Supplemental Table 8). Interestingly, genetic correlations with full-scale IQ were substantially higher for word reading (95% CI 0.70–0.85) than nonword reading (95% CI 0.50–0.68), likely reflecting the importance of reading skills for verbal tests of cognition. Genetic correlations of reading-/language-related traits with performance IQ (range 0.20–0.35) were much lower than those for full-scale IQ, and the 95% confidence intervals did not overlap. Indeed, only word reading showed a significant genetic correlation with performance IQ. Significant trait-specific genetic correlations were observed for components 2-4 of the Danish school grade decomposition analysis^18^. Component 2, reflecting relatively better school grade performance in language than mathematics (as compared to peers), was positively correlated with both GenLang reading traits. Component 3, reflecting relatively better school grade performance in oral than in written exams, showed significant negative correlations with phoneme awareness and spelling. Lastly, component 4, reflecting relatively better school grade performance in Danish than in English, showed significant negative correlation with nonword repetition. The non-cognitive abilities involved in educational attainment identified in the GWAS-by-subtraction study^17^ showed small but significant positive genetic correlations with word reading, nonword reading and nonword repetition.

Genomic structural equation modelling (GenomicSEM)^19^ was used to further investigate the shared genetic architecture of the five reading- and language-related traits, together with performance IQ, full-scale IQ and educational attainment (Supplemental Table 9). In the final model (Figure 1B), the first factor explains variation in nonword reading, spelling and phoneme awareness, word reading and full-scale IQ. For the first three of these traits, there is no evidence for additional genetic influences, suggesting high genetic similarity. The second factor explains additional variation in full-scale IQ, and is also related to performance IQ and educational attainment. The third factor explains variation in nonword repetition, word reading and educational attainment. Factors one and three are highly correlated, indicating that the genetic architecture underlying word reading does not differ much from nonword reading, spelling and phoneme awareness. Nonword repetition, on the other hand, is genetically more distinct, as indicated by evidence for specific genetic influences not captured by the model. Specific genetic influences were also evident for full-scale IQ and educational attainment. Thus, although reading- and language-related traits show genetic overlaps with full-scale IQ and educational attainment, the model indicates that these traits also have unique unshared components, in line with genetic correlation estimates that are lower than 1.

### Genes and SNPs previously associated with reading-/language-related traits and disorders

Previous GWAS studies on reading-/language-related traits and disorders (Supplemental Table 10) have identified very few associations exceeding genome-wide significance. In those prior studies, there were a total of 48 independent SNPs meeting a less stringent threshold of p<1×10^−6^ in the respective GWAS. We ran look-ups of each of those SNPs in our GenLang GWAS meta-analysis results (Supplemental Table 11). Where SNP associations passed a threshold adjusted for multiple testing of 48 SNPs and 2.15 independent GenLang traits (p<4.84×10^−4^), we then reran the association analyses after exclusion of the original cohort(s) in which the association was first identified, to evaluate independent effects beyond those of the earlier study. According to these criteria, only one SNP, rs1555839, previously associated with rapid automatized naming in the GRaD cohort^5^, yielded a significant signal in the remainder of the GenLang cohorts, showing association with spelling (p=3.33×10^−4^). Rs1555839 is one of five SNPs that reached the threshold for genome-wide significance in the original GWAS study of the GRaD cohort.

Some 20 genes have been described as candidate genes for reading-/language-related traits and disorders, based on a range of different mapping approaches^2^. Supplemental Table 12 gives gene-based p-values from our GenLang GWAS meta-analysis, calculated by MAGMA^20^, for each of these genes. Variation in one of these genes, namely *DCDC2*, showed association with nonword reading that passed the significance threshold for multiple testing of 20 genes and 2.15 independent GenLang traits (p<0.0012). *DCDC2* was originally identified in a linkage region for dyslexia susceptibility, and SNPs in and near this gene were subsequently associated with dyslexia in candidate gene studies^21^, although some investigations including a meta-analysis of seven studies have failed to support this^22^. Many dyslexic children perform poorly on nonword reading tests^23^. No single candidate SNP highlighted in prior studies of *DCDC2*, nor in any other candidate gene, was significantly associated with any of the traits in the GWAS meta-analysis results after correction for 54 SNPs and 2.15 independent GenLang traits (p<4.31×10^−4^) (Supplemental Table 13).

### Multivariate GWAS analysis for word reading

To improve the power of our GWAS meta-analysis and to gain insights into biological pathways shared by the traits of interest, utilizing the high genetic correlations between the traits, a multivariate GWAS analysis was performed with MTAG^24^. This approach improves the effect estimates of univariate meta-analysis results by incorporating information of other genetically correlated traits. Although MTAG generates output for each input trait, the results for word reading were used for all follow-up analyses, because the univariate word reading GWAS meta-analysis has the largest sample size (27,180 for the European-ancestry analysis, compared to 12,411-17,278 for the other three traits). Although no single SNP reached genome-wide significance in this multivariate word reading analysis (Supplemental Figure 4), the SNP-based heritability increased from 0.16 (se=0.04) to 0.29 (se=0.02). MTAG estimated the GWAS equivalent sample size of the multivariate word reading results as 41,783, compared to 27,180 for the univariate results, further indicating the increased power provided by MTAG. The multivariate word reading results were used for an array of follow-up analyses utilizing genetic correlations, gene property analysis in MAGMA and partitioning heritability.

### Genetic correlation with structural brain imaging traits

We performed a literature review to select 58 structural neuroimaging traits in the UK Biobank encompassing brain regions and white matter tracts with known links to aspects of reading and language (see Methods). These traits included surface based morphometry (SBM; surface area and thickness; Supplemental Figure 6) phenotypes and diffusion tensor imaging (DTI; mean and weighted mean fractional anisotropy; Supplemental Figure 7) results. As many of these brain-based phenotypes were significantly correlated with each other, genetic correlations among their summary statistics were calculated (Supplemental Figure 8) and were used to calculate the number of independent traits for multiple testing correction. One trait showed significant genetic correlation with the multivariate word reading results (p<0.05 / 24.85 independent traits = 2.01×10^−3^): the surface area of the banks of the superior temporal sulcus (STS) of the left hemisphere (Figure 2; Supplemental Table 14). Functional MRI studies have linked this region to different aspects of written and spoken language processing^25-28^. To investigate further how the different reading- and language-related traits contribute to these findings, we went on to specifically assess genetic correlations for banks of the left STS with the results from the original univariate GenLang GWAS meta-analyses, finding consistent correlations (range 0.18–0.23, Supplemental Table 15).

**Figure 2:**
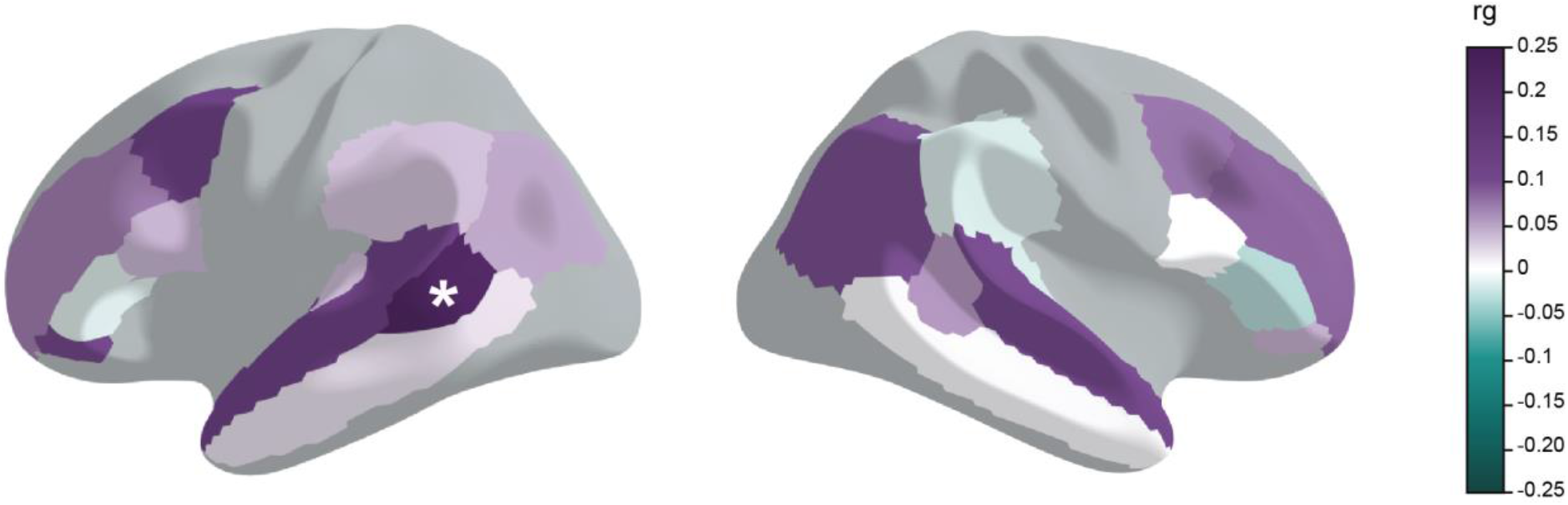
The multivariate word reading results show significant genetic correlation with the cortical surface area around the left superior temporal sulcus. Genetic correlations (rg) were estimated with LDSC. Included traits are 58 structural brain imaging traits from the UK Biobank selected based on known links of regions and circuits with language processing. The results of the 22 cortical surface areas are shown; grey areas were not included in the analysis. *Significant genetic correlation after correcting for 24.85 independent brain imaging traits (p<2.01×10^−3^). Full results can be found in Supplemental Table 14 and Supplemental Figure 6 and 7.

### Genetic correlation with traits from the UK biobank and brain-related traits from LD hub

We went on to assess genetic correlations of the multivariate word reading results with 20 cognitive, education, neurological, psychiatric and sleeping-related traits and 515 additional UK Biobank traits using LD hub. To further investigate overlaps with IQ, genetic correlations between these 535 traits and the published GWAS summary statistics for full-scale IQ^16^ (n=257,828) were obtained as well. A total of 143 traits showed significant genetic correlations with the multivariate word reading results after correction for multiple testing (p<0.05/(535*2)=4.67×10^−5^; Supplemental Table 16), while 245 traits were genetically correlated with full-scale IQ; 135 traits showed significant correlations both with word reading and full-scale IQ. Traits with strong genetic correlations with either word reading, full-scale IQ or both traits were related to education, eyesight, chronotype, wellbeing, lifestyle, physical health and exercise and socioeconomic status. Representative traits are plotted in Figure 3. The genetic correlations of reading and full-scale IQ with other traits had similar directions and effect sizes, but several differences were evident. Cognitive and education-related traits were more highly genetically correlated with full-scale IQ than reading. In addition, several psychiatric and wellbeing traits showed significant (negative) genetic correlations with full-scale IQ but not with reading, for example depressive symptoms, cross-disorder susceptibility (from the Psychiatric Genomics Consortium GWAS), and tense, hurt and nervous feelings. In contrast, several traits related to physical health and lifestyle showed larger genetic correlations with the multivariate reading results than with full-scale IQ, including BMI, reduced alcohol intake as a health precaution, and usual walking pace.

**Figure 3:**
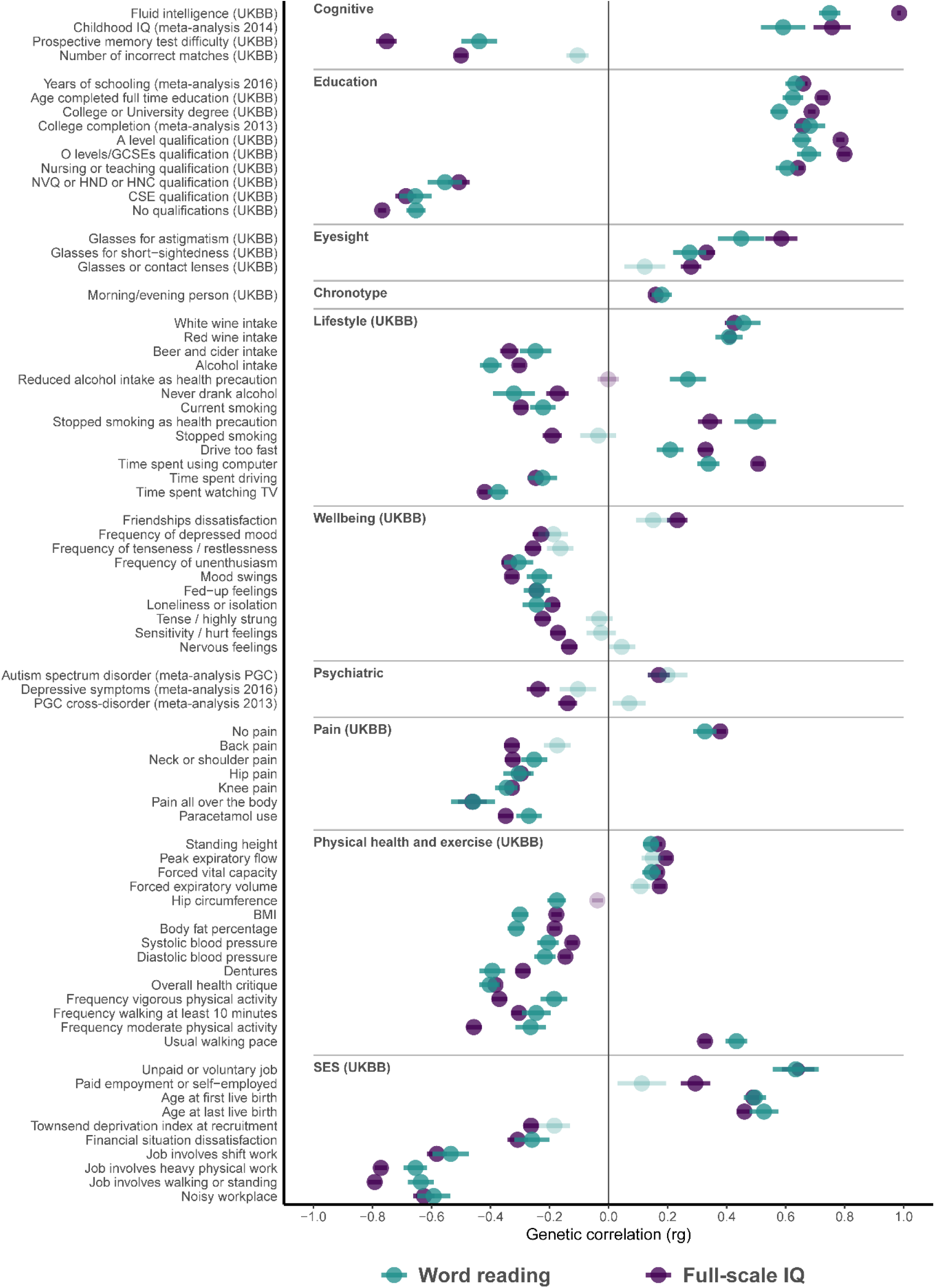
Genetic correlation results of the multivariate reading analysis, with comparisons to those for the largest published GWAS of cognitive performance in LDhub. Summary statistics for full-scale IQ (n=257,828) were obtained from the Social Science Genetic Association Consortium^16^. Genetic correlations between the multivariate word reading results (blue-green), full-scale IQ (purple) and traits in LDhub reveal an overlap with cognitive traits, education, eyesight, chronotype, lifestyle, wellbeing, psychiatric disorders, pain, physical health and exercise, and socioeconomic status. A subset of representative traits is shown, out of 143 traits that showed significant associations with the multivariate word reading results and 245 traits that showed significant correlations with full-scale IQ, of which 135 traits overlap, after correction for multiple testing for 535*2 traits (p<4.67 × 10^−5^). Significant correlations are shown in dark colours, non-significant correlations in light colours. Full results can be found in Supplemental Table 16. Genetic correlation (rg) is presented as a dot and error bars indicate the standard error.

### Evolutionary analysis

We used LDSC heritability partitioning^29^ to study the heritability enrichment of the multivariate reading results for five annotations reflecting different aspects of human evolution spanning periods from 30 million to 50,000 years ago (Figure 4; Supplemental notes). Annotations include human gained enhancers (HGE) active in fetal and adult brain tissue, ancient selective sweep regions, Neanderthal introgressed SNPs and Neanderthal depleted regions. The multivariate reading results were significantly enriched for Neanderthal depleted regions, after correction for the five annotations analysed (p<0.01) (Supplemental Table 17). These regions are large stretches in the human genome that are depleted for Neanderthal ancestry, possibly due to critical functions in *Homo sapiens* and intolerance to gene flow^30^.

**Figure 4:**
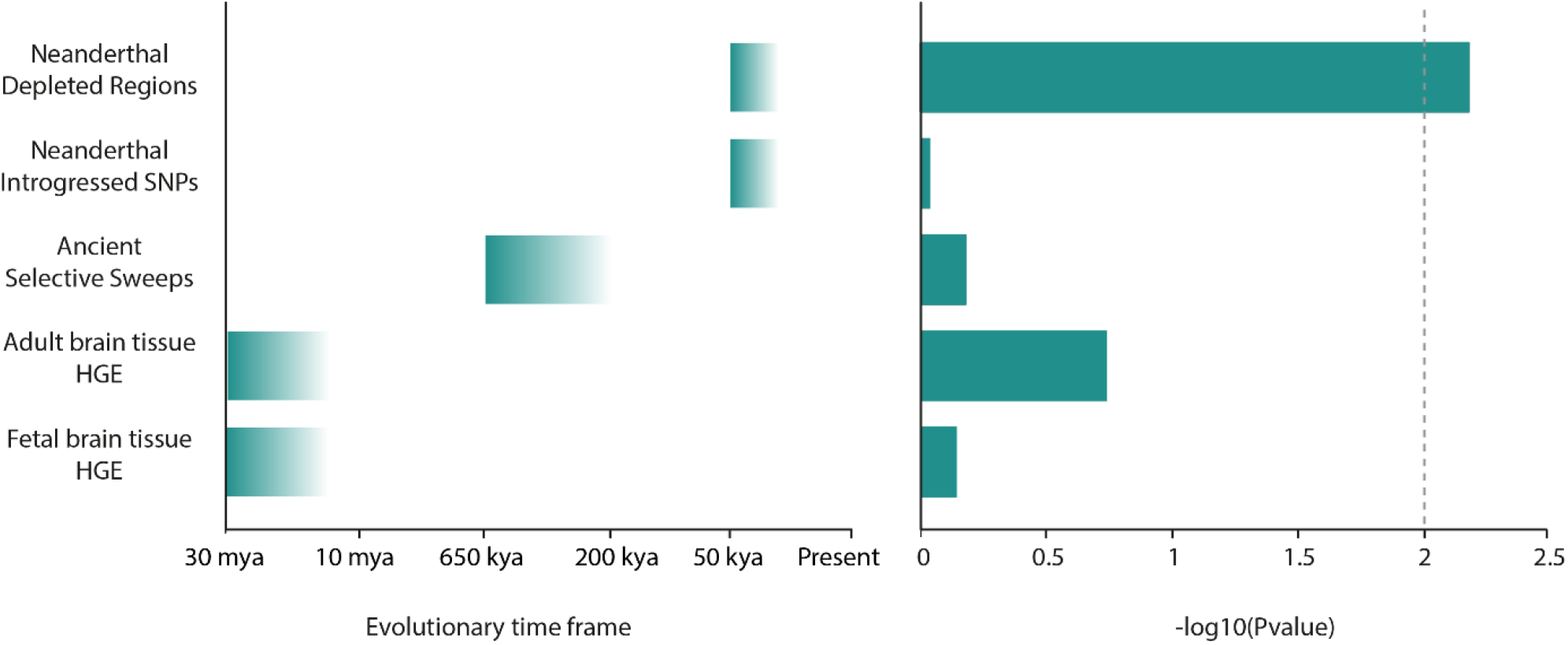
LDSC heritability partitioning of multivariate reading GWAS identifies significant enrichment in Neanderthal depleted regions. Five annotations were studied, reflecting different aspects of human evolution from diverse periods from 30 million to 50,000 years ago. Left: Overview of the approximate time frames captured by the five annotations. Mya: million years ago; kya: thousand years ago; HGE: human gained enhancers. Right: The -log-10 p-values of the partitioning heritability analysis of the multivariate reading results. The dashed line shows the p-value threshold for significant enrichment after Bonferroni-correction for testing 5 annotations. Results are also available in Supplemental Table 17.

### Functional enrichment using heritability partitioning and MAGMA gene property analysis

We applied LDSC heritability partitioning analyses^29^ to determine whether the heritability of the multivariate reading analysis is enriched in specific functional regions of the genome. We used 489 annotations reflecting tissue-specific chromatin signatures, as most variants identified in GWAS studies are located outside coding regions and are often found enriched in functional regions of the genome such as promoters, enhancers and regions with open chromatin. After correcting for testing 489 annotations (p<1.02×10^−4^), three annotations showed significant heritability enrichment for reading: histone-3 lysine-4 monomethylation (H3K4me1) in two fetal brain samples and the adult brain germinal matrix (Figure 5; Supplemental Figure 9 and Table 18). H3K4me1 is considered a marker for enhancer regions.

**Figure 5:**
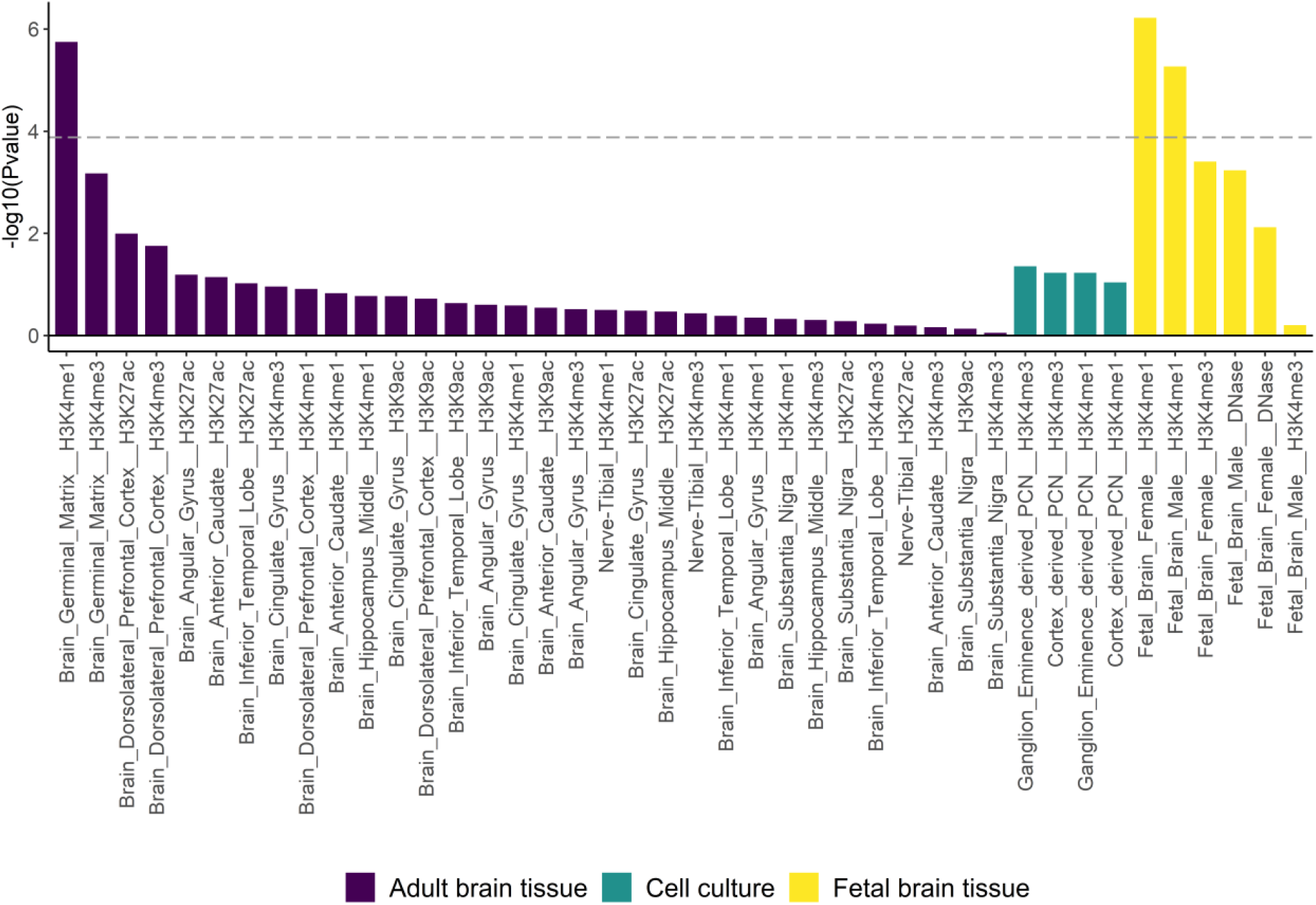
SNP-based heritability of reading is significantly enriched in brain enhancers. 489 annotations of tissue-specific chromatin signatures were used to analyse the multivariate reading results with LDSC heritability partitioning. Only brain annotations are shown; full results are available in Supplemental Figure 9 and Supplemental Table 18. PCN: primary cultured neurospheres. The graph shows log-10 p-values on the y-axis and brain region-specific chromatin mark on the x-axis. The dashed line shows the p-value threshold for significant enrichment after Bonferroni correction for testing 489 annotations.

Next, we used MAGMA gene property analysis^20^ to study whether the multivariate reading results are enriched in a specific tissue or brain cell type, using tissue-specific and cell type-specific gene expression data in FUMA^31,32^. As MAGMA corrects for average expression, each comparison can only answer the question of whether the tissue or cell type is more related to the multivariate reading results than the average of the tissues or cell types in the dataset. After correction for 83 tissues (p<6.02×10^−4^), no relation was found of the multivariate reading results with tissue-specific gene expression patterns of adult tissues from GTEx and brain tissues of a specific (developmental) time from Brainspan (Supplemental Figure 10 and Table 19). In the cell type-specific gene expression analysis, three single-cell RNA-sequencing datasets of embryonic, fetal and adult brain tissue were used. After correction for 142 cell types (p<3.09×10^−4^), the multivariate reading results were significantly associated with one of the mature neurons from the fetal dataset: red nucleus neurons (Supplemental Figure 11 and Table 20). This association could reflect an association with this specific nucleus, or with the higher maturity of the neurons compared to the other cell types in the fetal dataset. The red nucleus is a large subcortical structure in the ventral midbrain that is part of the olivocerebellar and cerebello-thalamo-cortical systems. It plays an important role in locomotion and non-motor behaviour in various animal species, and in humans might also play a role in higher cortical functions^33^. A trend was observed for a relation with GABAergic neurons from the embryonic dataset.

## Discussion

We performed GWAS meta-analysis of five quantitative reading- and language-related traits: word reading, nonword reading, spelling, phoneme awareness and nonword repetition, in sample sizes (up to ∼34,000 participants) that are substantially larger than any prior genetic analyses of reading and/or language skills assessed with neuropsychological tools (n=1,331-10,819)^3-5,34-36^. We identified genome-wide significant associations for word reading (rs11208009 at chromosome 1, p=1.10×10^−8^), highlighting *DOCK7, ATG4C, ANGPTL3* and *USP1* as potential new candidate genes. Other SNPs from the locus have been previously associated with triglyceride and cholesterol levels^37^, but may represent an independent association signal. Robust SNP-based heritabilities were observed, ranging from 0.13 for nonword repetition to 0.26 for nonword reading. These SNP-based heritabilities are similar to those of the related trait dyslexia (estimates range from 0.15 to 0.25 on a liability scale)^38,39^, psychiatric traits such as ADHD symptoms in adults (0.22)^40^, and brain imaging traits such as cortical surface area (range 0.12–0.33 for different cortex regions)^41^ and cortical thickness (range 0.08–0.26)^41^ and they are larger than that of psychiatric traits such as major depression (0.08)^42^ and alcohol dependence (0.09)^43^. So, despite highlighting the need for larger sample sizes to identify more than one genome-wide significant locus, our GenLang GWASs already allowed for multiple informative follow-up analyses based on the full dataset.

Word reading, nonword reading, spelling, and phoneme awareness largely share the same genetic architecture, as is evident from genetic correlation analyses, as well as the three-factor model in which one factor explains most of the variation in these four traits. Nonword repetition, IQ and educational attainment have, at least in part, a different genetic foundation, as reflected in the residual genetic variation of these traits not captured by the model. These findings are consistent with multiple behavioural studies showing a distinction between nonword repetition and other reading- and language-related traits^44^. They are also in line with results of recent structural equation modelling of genetic trait interrelatedness for 11 different reading- and language-related measures in the ALSPAC cohort, which identified a shared genetic factor accounting almost fully for the genetic variance in literacy-related phenotypes, but for only 53% of that in nonword repetition^45^.

The five reading- and language-related traits in GenLang also showed high genetic overlaps with full-scale IQ and educational attainment, in line with the widespread pleiotropy found between many aspects of cognitive functioning including language, reading, mathematics and general cognitive ability^4,46^. Given the evidence of a highly shared genetic architecture between word reading, nonword reading, spelling, and phoneme awareness, we performed a multivariate analysis with MTAG, to increase our power and reduce multiple-testing constraints for follow-up analyses of biological pathways. The multivariate word reading results showed genetic correlations with phenotypes related to education, eyesight, chronotype, wellbeing, lifestyle, physical health, exercise and socioeconomic status. These associations may in a large part reflect shared biology between the reading-/language-related traits and cognitive abilities, educational attainment and socioeconomic status. No significant genetic correlations were observed with neuropsychiatric disorders based on available data in LDhub. However, we note that summary statistics from the current investigation have also been used to investigate genetic overlaps with self-report of dyslexia diagnosis in an independent GWAS by 23andMe (∼52k cases), yielding substantial negative genetic correlations between GenLang quantitative traits and dyslexia status (e.g. -0.71 for word reading, -0.75 for spelling) as reported in a preprint by Doust et al. 38^39^. The pattern of findings is also likely to be influenced by genetic nurture^47^, relating to the socioeconomic status of the family, as was recently found for polygenic score analyses of cognitive traits^48^. Future investigations including information about nontransmitted alleles^49^, or data from siblings^48^, may help to disentangle pleiotropy from genetic nurture.

Beyond pleiotropy, some evidence of differences in trait aetiology was observed in our genetic correlation analysis with components (identified via decomposition analysis) from genome-wide analysis of school grades in the Danish iPSYCH cohort^18^ and with results of a GWAS-by-subtraction analysis of educational attainment and cognitive performance^17^. The Danish school grade “component 3”, corresponding to relatively better performance in oral than written exams, showed negative genetic correlations with phoneme awareness and spelling results in GenLang. In other words, higher scores on phoneme awareness and spelling appear to be genetically correlated with better performance in written than in oral exams. This may reflect the greater importance of phoneme awareness and spelling for proficient writing than for oral language. “Component 4”, corresponding to relatively better performance in Danish (the native language of the participants in that study) as compared to performance in English, showed negative genetic correlations with our GenLang nonword repetition measure, possibly reflecting the particular importance of verbal short-term memory in second-language learning^50,51^. The results of the GWAS-by-subtraction, previously suggested to represent so-called “non-cognitive” abilities related to educational attainment such as motivation, curiosity and persistence, were genetically correlated with word reading, nonword reading, and nonword repetition in GenLang.

Human abilities to process spoken and written language depend on an array of distributed brain circuits^52-56^. We performed a genetic correlation analysis of our multivariate GenLang GWAS with summary statistics from 58 MRI-based neuroanatomical phenotypes, chosen because they concerned brain areas and/or tracts with known links to language processing^53-56^. This selection included brain regions involved in early modality-specific pre-processing of spoken and written language, for example the auditory regions in transverse temporal gyrus, also known as Heschl’s gyrus, and the superior temporal gyrus (spoken language), and several other temporal regions (written language). Other regions included are involved in aspects of language comprehension, including the middle temporal gyrus, the inferior parietal area and parts of the inferior frontal gyrus. We identified a significant genetic correlation of our multivariate GWAS with cortical surface area of the banks of the superior temporal sulcus (STS) on the left hemisphere. The STS is a location where the processing of spoken and written language converges, in between modality-specific pre-processing and language comprehension^25-28^. A broad range of language-related functions have been previously linked with the left STS through (meta-analyses of) fMRI and PET studies, including those essential for the reading- and language-related traits included in the GWAS meta-analyses: sublexical processing of speech^57,58^ and representation of phonological word forms^59^. The importance of this brain area for reading-related traits is also evident from a meta-analysis of structural MRI studies that found lower grey matter volume in the left STS related to reading disability and poor reading comprehension^60^. Thus, findings from the genetic correlation analysis are consistent with the role of the STS as a hub where the processing of different language modalities gets integrated, as well as the lateralization of such functions.

Capacities for acquiring spoken and written language appear to be unique to our species, involving underlying skills that emerged on the lineage leading to modern humans^61,62^. We used heritability partitioning to analyse five annotations representing different timeframes and aspects of human evolution. Genomic regions that are significantly depleted of Neanderthal ancestry were enriched for genetic variants showing associations in our multivariate GenLang GWAS. Neanderthal depleted regions are thought to correspond to genomic loci that were intolerant to the gene flow from Neanderthal populations into *Homo sapiens* which took place around 50-60 thousand years ago^30^. These loci are enriched for promoters and regions conserved in primates^63^, as well as enhancers present in many tissues and those specific for fetal brain and muscle^64^. Only brain enhancers show signs of stringent purifying selection against Neanderthal variation, indicating that these regions mark parts of the genome where variation has a high probability of deleterious consequence^64^. Our results may reflect divergent selection acting on skills that underlie language in humans compared to Neanderthals. However, in our analyses we could not pinpoint any specific timeframe of human evolution during which genetic variants associated with reading- and language-related traits were introduced.

Regarding functional implications of our findings, heritability partitioning analyses of the multivariate GenLang GWAS results identified enrichment in enhancer regions present in fetal brain tissue and the adult germinal matrix. The enhancer regions of the germinal matrix are highly similar to those of fetal brain tissues, and not the other adult brain tissues we studied, likely reflecting the neural stem cell population present in that tissue^65^. In these analyses there was no specific association with one particular brain cell type. However, follow-up work with MAGMA, using single-cell RNA sequencing data from fetal, embryonic and adult brain, uncovered a significant association with fetal neurons from the Red nucleus, which may relate to the more adult state of these neurons compared to the other cell types in the fetal dataset^66^. The MAGMA analysis could not be used to test for replication of the association with (fetal) brain tissue of the LDSC heritability partitioning, as results in this case are corrected for the association with the average expression of the dataset^20,31^, and the datasets with fetal data only included brain samples.

We used the GWAS meta-analysis results to investigate evidence for association of previously reported candidate SNPs/genes and suggestive genome-wide screening results from prior studies of reading-/language-related traits and disorders. Out of the 54 candidate SNPs and 20 candidate genes that we assessed (none of which met genome-wide significance), only *DCDC2* yielded an association that survived correction for multiple testing in the context of targeted replications. This locus showed association only at the gene-based level and with one trait: nonword reading. Some previously reported associations in the literature could reflect the specific language, phenotype, or recruitment procedure of the cohort in which the gene or variant was investigated, and/or differences between contributions of common and rare variation at a locus of interest. Yet the lack of support here also suggests that false positive results have made an impact on the field, most likely related to limited sample size in prior reports, which is known to elevate the risk of type-1 error^67^. Few SNPs have shown genome-wide significant associations (p<5 × 10^−8^) in previous GWAS studies of quantitative reading-/language-related traits^3,5^. In our GWAS-meta analysis, the SNP rs1555839, previously associated with rapid automatized naming and rapid alternating stimulus^5^, was significantly associated with spelling. Overall, these results highlight the need for a genome-wide perspective, and the importance of large well-powered samples, if we are to obtain reliable insights into the role of common genetic variants in language- and reading-related traits.

Reading- and language-related phenotypes pose special challenges for scaling-up genetic analysis, since psychometric assessments can be labour-intensive to administer and score, and because of the heterogeneity introduced by differences in assessment tools, ages, populations and languages, among other factors^68,69^. One-item questions have enabled increases in sample size for GWAS of a wide range of traits and disorders, especially when available through large resources such as the UK Biobank. For example, a recent paper reporting development of a polygenic index repository included a polygenic score for a one-item question asking adult participants (customers of 23andMe) “at what age did you start to read?”^70^. This “age-started-reading” item was somewhat confusingly referred to in that study as “childhood reading”. While answers to the question were available in a large sample (n of >173k), the item has substantial limitations, including its reliance on adult self-report of the timing of specific events from early childhood, ambiguities over how to interpret the phrase “start to read”, as well as confounding effects of regional, cultural and historical differences in the age at which children first receive formal reading instruction. Moreover, the age a person started to read is a poor proxy of reading skill because of the large variation in the developmental trajectories of reading acquisition; in other words, children who learn to read relatively late in childhood can still become perfectly proficient readers. No validated questions have yet been described that adequately capture interindividual variability in reading and language skills in the normal range, which still requires administration of psychometric tests. The GenLang Consortium was established as an international effort by multiple research teams with the aim of overcoming such difficulties through a range of strategies, and enabling large-scale well-powered investigations of genomic underpinnings of these important traits.

This first wave of analysis from GenLang represents the largest GWAS meta-analyses for direct quantitative assessments of reading- and language-related abilities to date, including 22 cohorts with data available for at least one of the phenotypes. Nonetheless, although substantially increased over prior work in this area, sample sizes may still be considered relatively modest compared to the state-of-the-art for genetic association analyses of other complex traits. While they captured a significant proportion of the genetic variation underlying each phenotype, yielding several novel insights into the associated biology, detection of genome-wide significant loci was still limited. In addition, a number of phenotypes of interest (such as those that tap into syntactic skills) could not (yet) be pursued due inadequate sample sizes, even when combining data available from multiple cohorts. We note that despite our best efforts at harmonizing the included datasets, and limited signs of heterogeneity in the results based on Cochran Q statistics, LDSC intercepts and genetic correlations between subsets of the data, we cannot fully exclude that heterogeneity is introduced by the inclusion of data from different assessment tools, languages, and ages. The choice of assessment tools for future collection of reading- and language-related phenotypes for genomic studies, to increase the sample sizes of these GWAS meta-analyses and also to collect additional language-related phenotypes, should therefore be based at least partially on optimal matching with existing data. At the same time, we should invest in facilitating and simplifying the collection of language-related phenotypes, in part by developing and optimizing test batteries that could be reliably administered online in web/app-based settings. Indeed, these are major areas of focus for the GenLang consortium moving forward.

In summary, our GWAS meta-analyses of five reading- and language-related phenotypes in sample sizes of up to ∼34,000 participants have demonstrated significant SNP heritability for all traits, and identified genome-wide significant associations with word reading on chromosome 1 (rs11208009, p=1.098 × 10^−8^). Structural equation models revealed a single factor accounting for much of the genetic architecture underlying word reading, nonword reading, spelling and phoneme awareness, prompting a multivariate GWAS analysis of these four highly correlated traits. The multivariate results were genetically correlated with cortical surface area of the banks of the left STS, a brain region where the processing of spoken and written language comes together. Finally, partitioned heritability analyses showed enrichments in fetal brain enhancers, highlighting links to early brain development, and in Neanderthal depleted regions, suggesting that genomic regions associated with emerging language-related skills in *Homo sapiens* may have been intolerant to gene flow from other archaic hominins. These efforts by GenLang open up novel avenues for deciphering the biological underpinnings of spoken and written language.

## Material and methods

### Phenotypes

Word reading accuracy was measured as the number of correct words read aloud from a list in a time restricted or unrestricted fashion. Nonword reading accuracy was measured as the number of nonwords read aloud correctly from a list in a time restricted or unrestricted fashion. A nonword is a group of phonemes that looks or sounds like a word, obeys the phonotactic rules of the language, but has no meaning. Spelling accuracy was measured by the number of words correctly spelled orally or in writing, after being dictated as single words or in a sentence. Phoneme awareness was measured in phoneme deletion/elision and spoonerism tasks. Nonword repetition accuracy was measured by the number of nonwords or phonemes repeated aloud correctly. Performance IQ, included for follow-up analyses of the results of the reading- and language-related traits, employing a matching study design, was measured with nonverbal subtests of broader batteries that test for cognitive skills. The Supplemental Notes contain further details of phenotyping procedures, and Supplemental Table 1 provides an overview of the tests used to measure these phenotypes for each cohort.

### Study cohorts

The meta-analyses included GWAS summary statistics from 22 independent cohorts. These were, alphabetically, the Adolescent Brain Cognitive Development^SM^ Study (ABCD Study), the Avon Longitudinal Study of Parents and Children (ALSPAC), the ASTON cohort, the Brisbane Adolescent Twin Sample (BATS), the Basque Center on Cognition, Brain and Language (BCBL) cohort, the Colorado Learning Disabilities Research Center (CLDRC) cohort, the Early Language in Victoria Study (ELVS), the Familial Influences on Literacy Abilities (FIOLA) project, Generation R (GENR), the Genes, Reading, and Dyslexia (GRaD) study, the Iowa study, the NeuroDys cohort, the Netherlands Twin Register (NTR), the Pediatric Imaging, Neurocognition, and Genetics (PING) cohort, the Philadelphia Neurodevelopmental Cohort (PNC), the Raine Study, the SLI Consortium (SLIC) cohort, the Saguenay Youth Study (SYS), the Twins Early Development Study (TEDS), the Toronto cohort, the Oxford Dyslexia cohort (UKDYS), and the York cohort (see Supplemental Table 1 for demographic characteristics for each cohort). The cohorts were collected in different countries: in the USA, UK, the Netherlands, Australia, Canada, Spain, Austria, Germany, Switzerland, Finland, Hungary and France (ordered by sample size). Most participants are therefore from countries with English as their main language. Other languages spoken by participants are Dutch (n≤2,865, depending on trait), Spanish (n≤1,236), German (n≤1,227), Finnish (n≤323), French (n≤137) and Hungarian (n≤225). Most cohorts mainly include participants of European ancestry, with the exception of the GRaD cohort, which consists of individuals of African-American and Hispanic ancestry, and the ABCD Study, GenR and PING cohort, which are multi-ethnic. The sample sizes per cohort range from 104 to 10,187 participants (104 to 5,080 participants of European ancestry, defined by principal component analyses (PCA)). For each cohort, an Institutional Review Board (IRB) or ethical committee approved the respective studies, and participants provided informed consent.

Different measures for the reading- and language-related traits had been assessed in each cohort and could be included in the GWAS meta-analyses (see Supplemental Notes for details of each measure, and Supplemental Table 1 for an overview of the included measures and sample sizes for each cohort). Data from children, adolescents and young adults were included in the meta-analysis (age at time of assessment ranging from 5 years to 26 years). Outlier samples, based on the phenotype data (>4 SD), were removed for each phenotype separately. The phenotype data were then adjusted for covariates (age, age^2^, sex and ancestry principal components; age-normed phenotypes were not adjusted for age and age^2^). See Supplemental Table 2 for details on the included covariates per cohort. Phenotype data for word reading, nonword repetition and performance IQ were standardized to z-scores. Phenotype data for spelling, phoneme awareness and nonword reading were rank transformed to acquire normally distributed data for all cohorts. For follow-up analyses involving multiple phenotypes - the genetic correlation analysis with LDSC^71^ and the multivariate GWAS analysis with MTAG^24^ - a separate rank transformation was performed for word reading, nonword repetition and performance IQ to further harmonize the phenotype data processing. For male- and female-only association analyses, phenotype data were filtered, adjusted and transformed for male and female subsets separately.

The genotype data were subjected to stringent quality control according to a detailed analysis plan following standard procedures for GWAS, including SNP filters for minor allele frequency, call rate and Hardy-Weinberg equilibrium and sample filters including missingness and (for cohorts of unrelated individuals) relatedness. Cohort-specific details on quality control can be found in Supplemental Table 2. Individuals of European ancestry were identified using PCA-based analysis of genetic diversity. Individuals with non-European ancestry were excluded from all cohorts, with exception of ABCD, GenR, GRaD and PING. For the ABCD, GenR and PING cohorts, two association analyses were performed, one including and one excluding individuals of non-European ancestry. Results of datasets including individuals of non-European ancestry were excluded from follow-up analyses with LDSC^71^, MAGMA^20^ and MTAG^24^, since these methods utilize LD information based on European-ancestry reference data when raw genotyping data is not available (as is the case for this GWAS meta-analysis). The X chromosome was included by all cohorts except for NeuroDys. Genotype data were imputed using the Haplotype Reference Consortium version 1.1 panel for 20 out of the 22 cohorts, and using the 1000 Genomes Project Phase 3 reference panel for the GRaD and SYS cohorts. Single variant association analyses were performed using linear regression methods with the imputed additive genotype dosages for the full dataset, and for males and females separately. For the X chromosome, males were treated as homozygous diploids. Descriptions for each cohort of the samples, phenotype measures, genotyping, quality control and analysis procedures can be found in Supplemental Table 1 and 2.

### Meta-analyses

The summary statistics for each GWAS cohort for each trait were subjected to stringent quality control measures. SNPs were excluded from the meta-analyses based on low imputation quality scores <0.7, minor allele frequency <0.01 and minor allele count ≤10. Additional quality control of each summary statistics file was performed with EasyQC^72^.

Meta-analyses of the summary statistics were performed with METAL^73^ (version March 2011), with effect size estimates weighted using the inverse of the corresponding standard errors. A total of 13,633 to 33,959 individuals (12,411 to 27,180 individuals of PCA-selected European ancestry) of 10 to 19 cohorts (no trait was available from all 22 cohorts) were included in the GWAS meta-analyses for the different traits (Table 1 and Supplemental Table 1). SNPs for which data were available from less than 5,000 individuals were excluded from the meta-analysis results. For the heritability and genetic correlation analyses with LDSC, separate meta-analyses without genomic control correction were performed, because the LDSC regression intercept can be used to estimate a more powerful and accurate correction factor than genomic control^71^. Only data of individuals of the PCA-selected European subgroup were included, to allow use of pre-computed LD scores, as genotyping data of all cohorts was not available at a single site to allow the computation of LD-scores for the full partially admixed dataset.

To accommodate the multiple-testing burden present in performing separate meta-analyses for the five reading- and language-related traits, while taking into account the high phenotypic correlations between them, we calculated the effective number of independent variables (VeffLi) from the meta-analysis results using PhenoSpD^74^ (v1.0.0). The Bonferroni-corrected genome-wide significant P-value threshold was determined at 2.33×10^−8^ (5×10^−8^ / 2.15 independent traits).

We investigated the degree to which differences between cohorts in age distribution and phenotyping tools introduced heterogeneity in the meta-analysis results. First, Cochran’s Q test statistics, which assess whether estimated effect sizes are homogeneous across studies, were obtained with METAL, visualized with quantile-quantile plots (Supplemental Figure 3) and used to decide between a fixed-effects and random-effects meta-analysis. Based on these analyses, a fixed-effects meta-analysis was performed for all traits except for nonword repetition. Second, LDSC intercept and ratio were inspected to distinguish polygenicity from confounders^71^. Third, meta-analyses of subsets of the cohorts were performed, split up by mean age or the type of reading test applied. Heterogeneity caused by difference in mean age and type of reading test was studied by calculating genetic correlations between data subsets using LDSC^11^. In addition, meta-analyses for male- and female-only subsets of the data were run as sensitivity analyses, to investigate the degree to which males and females might show differences in SNP heritability for these traits and show genetic overlap as calculated with genetic correlation analyses.

### GenomicSEM

To investigate the high genetic correlations between the reading- and language-related traits, and with cognitive performance and educational attainment, we used genomic structural equation modeling (GenomicSEM; version 0.03)^19^ to model the joint genetic architecture. Summary statistics of the five GenLang traits and performance IQ, and published GWAS summary statistics for cognitive performance and educational attainment from the Social Science Genetic Association Consortium (https://www.thessgac.org/data)^16^ were used as input. GenomicSEM first runs multivariable LDSC to obtain genetic covariance and sampling covariance matrices. Next, exploratory factor analyses were run using a maximum-likelihood factor analysis, for models with one to four factors. Confirmatory factor analyses were then run in GenomicSEM for the exploratory model that explained the largest part of the variance in the data. To determine which factor loadings from the exploratory model to include, different models were fitted and compared using the following model fit indices: the p-value of the chi-square test, Akaike Information Criterion (AIC), Comparative Fit Index (CFI), and Standardized Root Mean Square Residual (SRMS). The model with the highest p-value, lowest AIC, CFI >0.9, and SRMR <0.1, was considered the best fitting model. (A p-value above 0.05 may not be possible when including summary statistics of large samples^19^).

### Multivariate GWAS analysis

A multivariate GWAS was performed on the four most highly correlated traits: word reading, nonword reading, spelling and phoneme awareness, using Multi Trait Analysis of GWAS (MTAG, v1.0.8)^24^, to maximize information for follow-up analyses on biological pathways, evolutionary significance, and so on. MTAG can perform a multivariate GWAS using summary statistics of different but related traits, while correcting for overlapping samples. Because MTAG takes its sample overlap estimates from LDSC^71^, univariate meta-analysis results including only individuals of the PCA-selected European subgroup were used. MTAG outputs a result for each trait; only the MTAG results for word reading were used, as word reading had the largest sample size, and because the MTAG results for the four traits were highly similar as a consequence of the high genetic correlation between the traits. For the follow-up analyses using LDSC^71^ and MAGMA^20^, the GWAS equivalent sample size estimated by MTAG, was used as sample size.

### Heritability and genetic correlation

LDSC^71^ (v1.0.0) was used to estimate genomic inflation and SNP-based heritability of the meta-analysis results, and to investigate genetic correlations^11^. All analyses were based on HapMap 3 SNPs only, and precalculated LD scores from the European 1000 Genomes reference cohort were used. For the LDSC analyses of the MTAG results, the GWAS equivalent sample size, estimated by MTAG, was used as sample size. The influence of confounding factors was tested by comparing the estimated intercept of LDSC to one, and the ratio of LDSC to zero. This ratio estimates the proportion of inflation in χ^2^ attributable to confounding, as opposed to true polygenic effects. SNP heritability was estimated based on the slope of the LDSC.

GWAS summary statistics for genetic correlation analyses with cognitive traits were obtained from the Social Science Genetic Association Consortium (full-scale IQ and educational attainment^16^; https://www.thessgac.org/data); the GWAS catalogue (noncognitive skills investigated with GWAS by subtraction^17^; ftp://ftp.ebi.ac.uk/pub/databases/gwas/summary_statistics/GCST90011874), and through collaboration with the iPSYCH consortium (GWAS analysis of Danish school grades^18^). PhenoSpD^74^ was used to calculate the effective number of independent variables (VeffLi) to inform the multiple testing correction. A total of 18.28 independent comparisons were performed in Figure 1, the p-value threshold is therefore set to p=2.74*10^−3^.

Publicly available GWAS summary statistics of neuroimaging traits were obtained via the Oxford Brain Imaging Genetics Server^75^ (http://big.stats.ox.ac.uk/). Out of 3,144 brain imaging-derived traits with summary statistics available from the UK Biobank, a total of 58 neuroanatomical phenotypes were selected based on their relevance to language processing. Brain imaging traits encompassed surface-based morphometric (SBM) and diffusion tensor imaging (DTI) phenotypes. For SBM, data were originally generated with Freesurfer by parcellation of the white surface (the surface area between the white and grey matter) using the Desikan-Killiany atlas. Both cortical surface area and mean cortical thickness were selected for brain areas that overlapped regions previously related to language processing, based on literature review^53-56^. For DTI, tracts spanning the extended language network^52^ were selected, and fractional anisotropy values derived from both tract-based-spatial statistics and probabilistic tractography were used (both mean and weighted-mean fractional anisotropy). Again, PhenoSpd was used to calculate the effective number of independent comparisons. A total of 24.85 independent brain imaging-derived traits were identified. Therefore the p-value threshold for a significant genetic correlation between the brain imaging-derived traits and the MTAG results was set to p=2.01×10^−3^ (0.05/24.85).In addition to our targeted analysis of brain imaging traits, genetic correlations were estimated between the MTAG results and summary statistics of 20 cognitive, education, neurological, psychiatric and sleeping-related traits and all 515 UK Biobank traits available in LD Hub^76^ (v1.9.3, http://ldsc.broadinstitute.org/ldhub/). These 535 traits comprise all phenotypes available through LDhub with relevance to brain function (beyond neuroimaging traits), and all available traits from the UK Biobank. Genetic correlations between these 535 traits and the published GWAS summary statistics for full-scale IQ^16^ were obtained as well. The Bonferroni corrected p-value threshold for significance of the LDhub results was 0.05 / (535*2) = 4.67×10^−5^. Genetic correlations may reflect pleiotropy, correlation between causal loci or spurious associations, and can inform about shared biological mechanisms and causal relationships between traits^77^.

### Functional Mapping and Annotation of GWAS meta-analysis results

The platform Functional Mapping and Annotation of Genome-Wide Association Studies^32^ (FUMA GWAS; https://fuma.ctglab.nl/; version 1.3.6a) was used to annotate the genome-wide significant variants and to calculate gene-based p-values. Using the SNP2GENE function, genome-wide significant loci were annotated with expression quantitative trait locus (eQTL) data from 4 different databases: GTEx V8 (brain samples only; http://www.gtexportal.org/home/datasets), the blood eQTL browser (http://genenetwork.nl/bloodeqtlbrowser/), the BIOS QTL browser (http://genenetwork.nl/biosqtlbrowser/), and BRAINEAC (http://www.braineac.org/). Loci were also annotated with information on previously associated traits from the GWAS catalog (https://www.ebi.ac.uk/gwas/).

### Gene and gene-set analysis

MAGMA^20^ (version 1.08) gene analysis in FUMA was used to calculate gene-based p-values from SNPs located in the body of the gene and in the region 1kb upstream to include SNPs located in the promoter region. Look-ups were performed for candidate genes proposed in prior literature on reading-, speech- and language-related traits and disorders. MAGMA accounts for gene-size, number of SNPs in a gene and LD between markers. The gene-based analysis was performed with default parameters (SNP-wide mean model), with the European 1000 Genomes reference cohort phase 3 as reference panel.

The MTAG results were further analysed using MAGMA gene property analyses, to study relationships with tissue-specific and cell type-specific gene expression patterns. Bulk RNA-sequencing data from GTEx V8 (http://www.gtexportal.org/home/datasets) and Brainspan (http://www.brainspan.org) of adult tissue samples and developmental brain samples were assessed in the SNP2GENE analysis in FUMA^32^. In addition, single-cell RNA-sequencing data from human embryonic (midbrain, 6-11 weeks post conception; Gene expression omnibus (GEO) accession number GSE76381), fetal (prefrontal cortex, 8-26 weeks post conception; GEO accession number GSE104276) and adult (Allen Brain Atlas cell types data of the middle temporal gyrus; http://celltypes.brain-map.org/) brain samples were used in the Cell Type analysis in FUMA^31^. MAGMA performs a one-sided test which essentially assesses the positive relationship between tissue specificity and genetic association of genes.

### Partitioning heritability of chromatin and evolutionary signatures

LDSC heritability partitioning^29^ was used to estimate the enrichment of heritability of the MTAG results in annotations reflecting tissue-specific chromatin modification patterns. Annotations were based on data from the Roadmap Epigenomics project and ENTEX, processed by Finucane et al.^78^.

In addition, LDSC heritability partitioning was used to study the association with several annotations reflecting evolutionary signatures and annotations from different periods along the lineage leading to modern humans, ranging from around 50,000 years ago back to 30 million years, adapting a pipeline recently published by Tilot et al.^79^. The following annotations reflecting evolutionary features were used (details in Supplemental notes): 1) adult brain Human Gained Enhancers^80^, 2) fetal brain Human Gained Enhancers^81^, 3) ancient selective sweep regions^82^, 4) Neanderthal-introgressed SNPs^83^, and 5) Neanderthal-depleted regions^30^. All enrichments are controlled for the baseline LD v2 model^29,84^, and heritability enrichment in human adult and fetal HGEs were additionally controlled for adult and fetal brain active regulatory elements.

## Data availability

The full GWAS summary statistics will be made freely available through the GWAS Catalog (https://www.ebi.ac.uk/gwas/) and the website of the GenLang network (www.genlang.org).

## Code availability

Code used to perform the meta-analysis and follow-up analyses is available at https://gitlab.gwdg.de/else.eising/genlang_quantitative_trait_gwasma.

## Author contributions

*Cohort PI*: AJOW, CLB, CEP, CH, DB, DIB, DFN, EvB, GSK, JJM, JRG, KL, LR, MC, MJW, MMPvdS, MMN, NGM, PFdJ, RKO, SR, SP, TP, ZP

*Genetics analysis*: AG, AGA, BSP, BMM, CAW, CYS, DFN, DTT, EE, ELdZ, EV, FA, GSK, GZ, JJH, JAHC, KR, KGW, KMP, LJS, MB, MvD, NMS, NHS, PRJ, SDG, SP, TK, TFMA, YF, ZL

*Phenotype data analysis*: AG, ATM, BSP, CAW, CYS, DB, EV, ELdZ, EvB, FR, GSK, JAHC, KGW, KL, KMP, KR, MB, ML, PFdJ, SR, TK, TCB, UM, ZL

*Genetics collection*: AM, ATM, CEP, CF, DB, EvB, FC, FR, GSK, JBTo, JCD, JJH, KL, MB, MC, MJW, MMN, NGM, SEF, SP, SR, TFMA, UM, YF

*Phenotype data collection*: ATM, CEP, CH, DB, ELdZ, EvB, FR, GSK, GTL, HL, HT, JBTo, JCD, KEW, KL, KM, MB, MC, MJW, MMPvdS, MW, MCJF, NGM, PSD, RKO, SLG, SR, TCB, UM

*Centrally done analysis*: EE

*Writing team*: BSP, CF, EE, SEF

SEF and CF conceived the study; BSP, CF, EE and SEF designed the study; EE and MG organized data gathering; EE, BM and GA performed analyses to interpret the results; ADB, DD and VM provided summary statistics. All authors contributed to and approved the final manuscript.

## Acknowledgements

BM, BMM, BSP, CF, EE, EV, GA, MvD and SEF are supported by the Max Planck Society. AG and TFMA were supported by the Munich Cluster for Systems Neurology (SyNergy), and AG was supported by Fondazione Umberto Veronesi. ATM is supported by the National Health and Medical Research Council of Australia (NHMRC) Fellowships (1105008; 1195955) and Centre of Research Excellence grant (1116976). AJOW is supported by an Investigator Grant from the NHMRC (1173896). BSP and MvD are supported by the Simons Foundation Autism Research Initiative grant (514787 to BSP). CYS works in the Medical Research Council Integrative Epidemiology Unit at the University of Bristol (MC_UU_00011/3). DIB acknowledges the Royal Netherlands Academy of Science Professor Award (PAH/6635). EGW is supported by NICHD P50 HD 27802. FR is supported by Agence Nationale de la Recherche (ANR-06-NEURO-019-01, ANR-17-EURE-0017 IEC, ANR-10-IDEX-0001-02 PSL, ANR-11-BSV4-014-01), European Commission (LSHM-CT-2005-018696). HT is supported by a VICI grant of the Netherlands Organization for Scientific Research (NWO) and The Netherlands Organisation for Health Research and Development (ZonMW) (016.VICI.170.200). JCDF was supported by the National Institute of Child Health and Human Development (NICHD) (P50 HD 27802). JJM, JBTo, and TK were supported by the National Institutes of Health (NIH) (R01 DC014489). KP was supported by the Hospital for Sick Children Research Training Program (Restracomp). KR is supported by a Sir Henry Wellcome Postdoctoral Fellowship. MJS is supported by a Wellcome Trust Programme grant (WT082032MA). SP and FA are supported by the Royal Society (UF150663; RGF\EA\180141). TB is supported by Institut Pasteur, the Bettencourt-Schueller Foundation, Université de Paris.

We are extremely grateful to all the children, twins, families and participants who took part and are taking part in the 22 cohorts whose data contributed to these GWAS meta-analyses. We also thank the staff working on the different cohorts, including volunteers, study coordinators, interviewers, teachers, nurses, research scientists, general practitioners, midwives, psychologists, psychometrists, computer and laboratory technicians and colleagues who assisted in the quality control and preparation of the imputed GWAS data and the pharmacies and hospitals that were involved.

We gratefully acknowledge iPSYCH for sharing their summary statistics. The iPSYCH team was supported by grants from the Lundbeck Foundation (R102-A9118, R155-2014-1724, and R248-2017-2003), the National Institute of Mental Health (NIMH, 1U01MH109514-01 to ADB) and the Universities and University Hospitals of Aarhus and Copenhagen. The Danish National Biobank resource was supported by the Novo Nordisk Foundation. High-performance computer capacity for handling and statistical analysis of iPSYCH data on the GenomeDK HPC facility was provided by the Center for Genomics and Personalized Medicine and the Centre for Integrative Sequencing, iSEQ, Aarhus University, Denmark (grant to ADB).

## Adolescent Brain Cognitive Development ^SM^ Study

Data used in the preparation of this article were obtained from the Adolescent Brain Cognitive Development^SM^ (ABCD) Study (https://abcdstudy.org), held in the NIMH Data Archive (NDA). This is a multisite, longitudinal study designed to recruit more than 10,000 children age 9-10 and follow them over 10 years into early adulthood. The ABCD Study® is supported by the NIH and additional federal partners under award numbers U01DA041048, U01DA050989, U01DA051016, U01DA041022, U01DA051018, U01DA051037, U01DA050987, U01DA041174, U01DA041106, U01DA041117, U01DA041028, U01DA041134, U01DA050988, U01DA051039, U01DA041156, U01DA041025, U01DA041120, U01DA051038, U01DA041148, U01DA041093, U01DA041089, U24DA041123, U24DA041147. A full list of supporters is available at https://abcdstudy.org/federal-partners.html. A listing of participating sites and a complete listing of the study investigators can be found at https://abcdstudy.org/consortium_members/. ABCD consortium investigators designed and implemented the study and/or provided data but did not necessarily participate in the analysis or writing of this report. This manuscript reflects the views of the authors and may not reflect the opinions or views of the NIH or ABCD consortium investigators. The ABCD data used in this report are summarized in this NDA study (https://doi.org/10.15154/1522876).

## Aston cohort

Data collection for the Aston Cohort has been supported by funding from the European Union Horizon 2020 Programme (641652), The Waterloo Foundation (797/17290), and with the invaluable support of the participants and their families. Data analyses were supported by the St Andrews Bioinformatics Unit which is funded by Wellcome Trust ISSF awards (105621/Z/14/Z and 204821/Z/16/Z).

## Avon Longitudinal Study of Parents and their Children (ALSPAC)

We are extremely grateful to all the families who took part in this study, the midwives for their help in recruiting them, and the whole ALSPAC team, which includes interviewers, computer and laboratory technicians, clerical workers, research scientists, volunteers, managers, receptionists and nurses. GWAS data were generated by Sample Logistics and Genotyping Facilities at Wellcome Sanger Institute and LabCorp (Laboratory Corporation of America) using support from 23andMe. The UK Medical Research Council and Wellcome (Grant ref: 217065/Z/19/Z) and the University of Bristol provide core support for ALSPAC. A comprehensive list of grants funding is available on the ALSPAC website (http://www.bristol.ac.uk/alspac/external/documents/grant-acknowledgements.pdf). The publication is the work of the authors and they will serve as guarantor for the ALSPAC contribution to this article.

## The Basque Center on Cognition, Brain and Language (BCBL)

Data collection was supported by the Basque Government through the BERC program, and by the Agencia Estatal de Investigación through BCBL Severo Ochoa excellence accreditation.

## Brisbane Adolescent Twins Study (BATS)

The research was supported by the Australian Research Council (A7960034, A79906588, A79801419, DP0212016 and DP0343921), with genotyping funded by the NHMRC (Medical Bioinformatics Genomics Proteomics Program, 389891).

## Colorado Learning Disabilities Research Center (CLDRC)

The CLDRC was supported by a NICHD grant P50 HD 27802.

## The Early Language in Victoria Study (ELVS)

Funding by the NHMRC grant number 436958 is gratefully appreciated.

## Familial Influences on Literacy Abilities (FIOLA)

The FIOLA cohort (PIs EvB and PFdJ) is supported by the University of Amsterdam, the MPI Nijmegen, and by fellowships awarded to EvB (NWO’s Rubicon 446-12-005 and VENI 451-15-017). We are grateful to the NEMO Science Museum.

## Generation R

The Generation R Study is conducted by the Erasmus Medical Center in close collaboration with School of Law and Faculty of Social Sciences of the Erasmus University Rotterdam, the Municipal Health Service Rotterdam area, the Rotterdam Homecare Foundation, and the Stichting Trombosedienst & Artsenlaboratorium Rijnmond (STAR[1]MDC), Rotterdam.

## The Genes, Reading and Dyslexia (GRaD) Study

The GRaD Study was funded by the generous support of the Manton Foundation, the NIH (P50-HD027802; K99-HD094902), and the Lambert Family.

## Neurodys

We would like to acknowledge our project partners Catherine Billard, Caroline Bogliotti, Vanessa Bongiovanni, Laure Bricout, Camille Chabernaud, Yves Chaix, Isabelle Comte-Gervais, Florence Delteil-Pinton, Jean-François Démonet, Fabien Fauchereau, Florence George, Christophe-Loïc Gérard, Guillaume Huguet, Stéphanie Iannuzzi, Marie Lageat, Marie-France Leheuzey, Marie-Thérèse Lenormand, Marion Liébert, Emilie Longeras, Emilie Racaud, Isabelle Soares-Boucaud, Sylviane Valdois, Nadège Villiermet, and Johannes Ziegler. This project was funded by the EU FP6 grant to the Neurodys consortium, the SNSF grant 32-108130 and Austrian Science Fund (18351-B02).

## Netherlands Twin Register (NTR)

Funding from the NWO and ZonMW is gratefully acknowledged: Twin family database for behavior genomics studies (NWO 480-04-004); Longitudinal data collection from teachers of Dutch twins and their siblings (481-08-011), Twin-family-study of individual differences in school achievement (NWO-FES, 056-32-010), Genotype-phenotype database for behavior genetic and genetic epidemiological studies (ZonMw Middelgroot 911-09-032); Genetic influences on stability and change in psychopathology from childhood to young adulthood (ZonMW 912-10-020), Why some children thrive (OCW_Gravity program –NWO-024.001.003), Netherlands Twin Registry Repository: researching the interplay between genome and environment (NWO-Groot 480-15-001/674); BBMRI –NL (184.021.007 and 184.033.111): Biobanking and Biomolecular Resources Research Infrastructure; Spinozapremie (NWO-56-464-14192); Amsterdam Public Health (APH) and Amsterdam Reproduction and Development (AR&D) Research Institutes; European Science Council Genetics of Mental Illness (ERC Advanced, 230374); Aggression in Children: Unraveling gene-environment interplay to inform Treatment and InterventiON strategies (ACTION) project (European Union Seventh Framework Program (FP7/2007-2013) no 602768); Rutgers University Cell and DNA Repository cooperative agreement (NIMH U24 MH068457-06); Developmental Study of Attention Problems in Young Twins (NIMH, RO1 MH58799-03); Grand Opportunity grant Developmental trajectories of psychopathology (NIMH 1RC2 MH089995) and the Avera Institute for Human Genetics.

## Pediatric Imaging, Neurocognition and Genetics Study (PING)

Data collection and sharing for this project was funded by the Pediatric Imaging, Neurocognition and Genetics Study (PING) (NIH grant RC2DA029475). PING is funded by the National Institute on Drug Abuse and the Eunice Kennedy Shriver NICHD. PING data are disseminated by the PING Coordinating Center at the Center for Human Development, University of California, San Diego.

## The Philadelphia Neurodevelopmental Cohort (PNC)

Support for the collection of the data sets was provided by grant RC2MH089983 awarded to Raquel Gur and RC2MH089924 awarded to Hakon Hakonarson. All subjects were recruited through the Center for Applied Genomics at The Children’s Hospital in Philadelphia. Reading performance was assessed as part of the cognitive test battery and collected by Gur et al.^85^. Genotyping was funded by an Institutional Development Award to the Center for Applied Genomics from The Children’s Hospital of Philadelphia and a donation from Adele and Daniel Kubert. We would also like to thank the NIH data repository. The dataset analysed was accessed via dbGaP number 12043 and 19848.

## The Raine Study

The Raine Study was supported by the NHMRC (572613, 403981 and 1059711), the Canadian Institutes of Health Research (MOP-82893), and WA Health, Government of Western Australia (Future Health WA G06302). Funding was also generously provided by Safe Work Australia. The authors gratefully acknowledge the NHMRC for their long-term funding to the study over the last 30 years and also the following institutes for providing funding for Core Management of the Raine Study: The University of Western Australia (UWA), Curtin University, Women and Infants Research Foundation, Telethon Kids Institute, Edith Cowan University, Murdoch University, The University of Notre Dame Australia and The Raine Medical Research Foundation. This work was supported by resources provided by the Pawsey Supercomputing Centre with funding from the Australian Government and Government of Western Australia.

## Saguenay Youth Study (SYS)

The Canadian Institutes of Health Research (ZP, TP), Heart and Stroke Foundation of Quebec (ZP), and the Canadian Foundation for Innovation (ZP) support the Saguenay Youth Study.

## SLI Consortium (SLIC)

SLI Consortium members are as follows: Wellcome Trust Centre for Human Genetics, Oxford: D. F. Newbury, N. H. Simpson, F. Ceroni, A. P. Monaco; Max Planck Institute for Psycholinguistics, Nijmegen: S. E. Fisher, C. Francks; Newcomen Centre, Evelina Children’s Hospital, St Thomas’ Hospital, London: G. Baird, V. Slonims; Child and Adolescent Psychiatry Department and Medical Research Council Centre for Social, Developmental, and Genetic Psychiatry, Institute of Psychiatry, London: P. F. Bolton; Medical Research Council Centre for Social, Developmental, and Genetic Psychiatry Institute of Psychiatry, London: E. Simonoff; Salvesen Mindroom Centre, Child Life & Health, School of Clinical Sciences, University of Edinburgh: A. O’Hare; Cell Biology & Genetics Research Centre, St. George’s University of London: J. Nasir; Queen’s Medical Research Institute, University of Edinburgh: J. Seckl; Department of Speech and Language Therapy, Royal Hospital for Sick Children, Edinburgh: H. Cowie; Speech and Hearing Sciences, Queen Margaret University: A. Clark, J. Watson; Department of Educational and Professional Studies, University of Strathclyde: W. Cohen; Department of Child Health, the University of Aberdeen: A. Everitt, E. R. Hennessy, D. Shaw, P. J. Helms; Audiology and Deafness, School of Psychological Sciences, University of Manchester: Z. Simkin, G. Conti-Ramsden; Department of Experimental Psychology, University of Oxford: D. V. M. Bishop; Biostatistics Department, Institute of Psychiatry, London: A. Pickles.

The collection and genetic characterisation of SLIC samples was funded by the Wellcome Trust (076566) and the UK Medical Research Council (G1000569).

## The Twins Early Development Study (TEDS)

TEDS is supported by a programme grant to RP from the UK Medical Research Council (MR/V012878/1 and previously MR/M021475/1), with additional support from the NIH (AG046938). The research leading to these results has also received funding from the European Research Council under the European Union’s Seventh Framework Programme (FP7/2007-2013/ grant agreement number 602768).

## Toronto

Support for this project was provided by grants from the Canadian Institutes of Health Research (MOP-133440).

## UK Dyslexia

Cohort recruitment and data collection was supported by Wellcome Trust (076566/Z/05/Z and 075491/Z/04) and The Waterloo Foundation (grants to JBTa and SP; 797–1720). Genotype data were generated and analyses funded by the EU (Neurodys, 018696) and the Royal Society (UF100463 grant to SP).

## York

The York cohort was funded by Wellcome Trust Programme Grant 082036/B/07/Z.

## References

1. Andreola, C. et al. The heritability of reading and reading-related neurocognitive components: A multi-level meta-analysis. Neurosci Biobehav Rev 121, 175–200 (2021).

2. Graham, S.A. & Fisher, S.E. Understanding Language from a Genomic Perspective. Annu Rev Genet 49, 131–60 (2015).

3. St Pourcain, B. et al. Common variation near ROBO2 is associated with expressive vocabulary in infancy. Nat Commun 5, 4831 (2014).

4. Gialluisi, A. et al. Genome-wide association scan identifies new variants associated with a cognitive predictor of dyslexia. Transl Psychiatry 9, 77 (2019).

5. Truong, D.T. et al. Multivariate genome-wide association study of rapid automatised naming and rapid alternating stimulus in Hispanic American and African-American youth. J Med Genet 56, 557–566 (2019).

6. Fisher, S.E. & Vernes, S.C. Genetics and the Language Sciences. Annual Review of Linguistics, Vol 1 1, 289–310 (2015).

7. Ehri, L.C. Learning to read words: Theory, findings, and issues. Scientific Studies of reading 9, 167–188 (2005).

8. Ehri, L.C. The development of spelling knowledge and its role in reading acquisition and reading disability. Journal of learning disabilities 22, 356–365 (1989).

9. Duncan, L.G. Language and Reading: the Role of Morpheme and Phoneme Awareness. Current Developmental Disorders Reports 5, 226–234 (2018).

10. Coady, J.A. & Evans, J.L. Uses and interpretations of non-word repetition tasks in children with and without specific language impairments (SLI). International journal of language & communication disorders 43, 1–40 (2008).

11. Bulik-Sullivan, B. et al. An atlas of genetic correlations across human diseases and traits. Nat Genet 47, 1236–41 (2015).

12. Watabe-Uchida, M., John, K.A., Janas, J.A., Newey, S.E. & Van Aelst, L. The Rac activator DOCK7 regulates neuronal polarity through local phosphorylation of stathmin/Op18. Neuron 51, 727–39 (2006).

13. Conklin, D. et al. Identification of a mammalian angiopoietin-related protein expressed specifically in liver. Genomics 62, 477–82 (1999).

14. Nijman, S.M. et al. The deubiquitinating enzyme USP1 regulates the Fanconi anemia pathway. Mol Cell 17, 331–9 (2005).

15. Zhang, L., Li, J., Ouyang, L., Liu, B. & Cheng, Y. Unraveling the roles of Atg4 proteases from autophagy modulation to targeted cancer therapy. Cancer Lett 373, 19–26 (2016).

16. Lee, J.J. et al. Gene discovery and polygenic prediction from a genome-wide association study of educational attainment in 1.1 million individuals. Nat Genet 50, 1112–1121 (2018).

17. Demange, P.A. et al. Investigating the genetic architecture of noncognitive skills using GWAS-by-subtraction. Nat Genet 53, 35–44 (2021).

18. Rajagopal, V.M. et al. Genome-wide association study of school grades identifies a genetic overlap between language ability, psychopathology and creativity. bioRxiv, 2020.05.09.075226 (2020).

19. Grotzinger, A.D. et al. Genomic structural equation modelling provides insights into the multivariate genetic architecture of complex traits. Nat Hum Behav 3, 513–525 (2019).

20. de Leeuw, C.A., Mooij, J.M., Heskes, T. & Posthuma, D. MAGMA: generalized gene-set analysis of GWAS data. PLoS Comput Biol 11, e1004219 (2015).

21. Meng, H. et al. DCDC2 is associated with reading disability and modulates neuronal development in the brain. Proc Natl Acad Sci U S A 102, 17053–8 (2005).

22. Zhong, R. et al. Meta-analysis of the association between DCDC2 polymorphisms and risk of dyslexia. Mol Neurobiol 47, 435–42 (2013).

23. Rack, J.P., Snowling, M.J. & Olson, R.K. The Nonword Reading Deficit in Developmental Dyslexia - a Review. Reading Research Quarterly 27, 28–53 (1992).

24. Turley, P. et al. Multi-trait analysis of genome-wide association summary statistics using MTAG. Nat Genet 50, 229–237 (2018).

25. Wilson, S.M., Bautista, A. & McCarron, A. Convergence of spoken and written language processing in the superior temporal sulcus. Neuroimage 171, 62–74 (2018).

26. Venezia, J.H. et al. Auditory, Visual and Audiovisual Speech Processing Streams in Superior Temporal Sulcus. Front Hum Neurosci 11, 174 (2017).

27. Friederici, A.D., Kotz, S.A., Scott, S.K. & Obleser, J. Disentangling syntax and intelligibility in auditory language comprehension. Hum Brain Mapp 31, 448–57 (2010).

28. Vigneau, M., Jobard, G., Mazoyer, B. & Tzourio-Mazoyer, N. Word and non-word reading: what role for the Visual Word Form Area? Neuroimage 27, 694–705 (2005).

29. Finucane, H.K. et al. Partitioning heritability by functional annotation using genome-wide association summary statistics. Nat Genet 47, 1228–35 (2015).

30. Vernot, B. et al. Excavating Neandertal and Denisovan DNA from the genomes of Melanesian individuals. Science 352, 235–9 (2016).

31. Watanabe, K., Umicevic Mirkov, M., de Leeuw, C.A., van den Heuvel, M.P. & Posthuma, D. Genetic mapping of cell type specificity for complex traits. Nat Commun 10, 3222 (2019).

32. Watanabe, K., Taskesen, E., van Bochoven, A. & Posthuma, D. Functional mapping and annotation of genetic associations with FUMA. Nat Commun 8, 1826 (2017).

33. Basile, G.A. et al. Red nucleus structure and function: from anatomy to clinical neurosciences. Brain Struct Funct 226, 69–91 (2021).

34. Luciano, M. et al. A genome-wide association study for reading and language abilities in two population cohorts. Genes Brain Behav 12, 645–52 (2013).

35. Harlaar, N., Hayiou-Thomas, M.E., Dale, P.S. & Plomin, R. Why do preschool language abilities correlate with later reading? A twin study. J Speech Lang Hear Res 51, 688–705 (2008).

36. Price, K.M. et al. Genome-wide association study of word reading: Overlap with risk genes for neurodevelopmental disorders. Genes Brain Behav 19, e12648 (2020).

37. Hoffmann, T.J. et al. A large electronic-health-record-based genome-wide study of serum lipids. Nat Genet 50, 401–413 (2018).

38. Gialluisi, A. et al. Genome-wide association study reveals new insights into the heritability and genetic correlates of developmental dyslexia. Mol Psychiatry (2020).

39. Doust, C. et al. Discovery of 42 Genome-Wide Significant Loci Associated with Dyslexia. medRxiv, 2021.08.20.21262334 (2021).

40. Demontis, D. et al. Discovery of the first genome-wide significant risk loci for attention deficit/hyperactivity disorder. Nat Genet 51, 63–75 (2019).

41. Grasby, K.L. et al. The genetic architecture of the human cerebral cortex. Science 367(2020).

42. Wray, N.R. et al. Genome-wide association analyses identify 44 risk variants and refine the genetic architecture of major depression. Nat Genet 50, 668–681 (2018).

43. Walters, R.K. et al. Transancestral GWAS of alcohol dependence reveals common genetic underpinnings with psychiatric disorders. Nature neuroscience 21, 1656–1669 (2018).

44. Bishop, D.V. Developmental cognitive genetics: how psychology can inform genetics and vice versa. Q J Exp Psychol (Hove) 59, 1153–68 (2006).

45. Shapland, C.Y. et al. Multivariate genome-wide covariance analyses of literacy, language and working memory skills reveal distinct etiologies. NPJ Sci Learn 6, 23 (2021).

46. Kovas, Y. & Plomin, R. Generalist genes: implications for the cognitive sciences. Trends Cogn Sci 10, 198–203 (2006).

47. Hart, S.A., Little, C. & van Bergen, E. Nurture might be nature: cautionary tales and proposed solutions. NPJ Sci Learn 6, 2 (2021).

48. Selzam, S. et al. Comparing Within- and Between-Family Polygenic Score Prediction. Am J Hum Genet 105, 351–363 (2019).

49. Kong, A. et al. The nature of nurture: Effects of parental genotypes. Science 359, 424–428 (2018).

50. Adlof, S.M. & Patten, H. Nonword Repetition and Vocabulary Knowledge as Predictors of Children’s Phonological and Semantic Word Learning. J Speech Lang Hear Res 60, 682–693 (2017).

51. Soroli, E., Szenkovits, G. & Ramus, F. Exploring dyslexics’ phonological deficit III: foreign speech perception and production. Dyslexia 16, 318–40 (2010).

52. Catani, S.J.F.a.M. Diffusion Imaging Methods in Language Sciences. in The Oxford Handbook of Neurolinguistics (ed. Schiller, G.I.d.Z.a.N.O.) (Oxford University Press, 2019).

53. Roehrich-Gascon, D., Small, S.L. & Tremblay, P. Structural correlates of spoken language abilities: A surface-based region-of interest morphometry study. Brain Lang 149, 46–54 (2015).

54. Perdue, M.V., Mednick, J., Pugh, K.R. & Landi, N. Gray Matter Structure Is Associated with Reading Skill in Typically Developing Young Readers. Cereb Cortex 30, 5449–5459 (2020).

55. Richardson, F.M. & Price, C.J. Structural MRI studies of language function in the undamaged brain. Brain Struct Funct 213, 511–23 (2009).

56. Price, C.J. The anatomy of language: a review of 100 fMRI studies published in 2009. Ann N Y Acad Sci 1191, 62–88 (2010).

57. Liebenthal, E., Desai, R.H., Humphries, C., Sabri, M. & Desai, A. The functional organization of the left STS: a large scale meta-analysis of PET and fMRI studies of healthy adults. Front Neurosci 8, 289 (2014).

58. Turkeltaub, P.E. & Coslett, H.B. Localization of sublexical speech perception components. Brain Lang 114, 1–15 (2010).

59. Okada, K. & Hickok, G. Identification of lexical-phonological networks in the superior temporal sulcus using functional magnetic resonance imaging. Neuroreport 17, 1293–6 (2006).

60. Eckert, M.A., Berninger, V.W., Vaden, K.I., Jr., Gebregziabher, M. & Tsu, L. Gray Matter Features of Reading Disability: A Combined Meta-Analytic and Direct Analysis Approach(1,2,3,4). eNeuro 3(2016).

61. Fitch, W.T. Empirical approaches to the study of language evolution. Psychon Bull Rev 24, 3–33 (2017).

62. Corballis, M.C. Language Evolution: A Changing Perspective. Trends Cogn Sci 21, 229–236 (2017).

63. Petr, M., Paabo, S., Kelso, J. & Vernot, B. Limits of long-term selection against Neandertal introgression. Proc Natl Acad Sci U S A 116, 1639–1644 (2019).

64. Telis, N., Aguilar, R. & Harris, K. Selection against archaic hominin genetic variation in regulatory regions. Nat Ecol Evol 4, 1558–1566 (2020).

65. Roadmap Epigenomics, C. et al. Integrative analysis of 111 reference human epigenomes. Nature 518, 317–30 (2015).

66. Zhong, S. et al. A single-cell RNA-seq survey of the developmental landscape of the human prefrontal cortex. Nature 555, 524–528 (2018).

67. Button, K.S. et al. Power failure: why small sample size undermines the reliability of neuroscience. Nat Rev Neurosci 14, 365–76 (2013).

68. Newbury, D.F., Monaco, A.P. & Paracchini, S. Reading and language disorders: the importance of both quantity and quality. Genes (Basel) 5, 285–309 (2014).

69. Deriziotis, P. & Fisher, S.E. Speech and Language: Translating the Genome. Trends Genet 33, 642–656 (2017).

70. Becker, J. et al. Resource profile and user guide of the Polygenic Index Repository. Nature Human Behaviour (2021).

71. Bulik-Sullivan, B.K. et al. LD Score regression distinguishes confounding from polygenicity in genome-wide association studies. Nat Genet 47, 291–5 (2015).

72. Winkler, T.W. et al. Quality control and conduct of genome-wide association meta-analyses. Nat Protoc 9, 1192–212 (2014).

73. Willer, C.J., Li, Y. & Abecasis, G.R. METAL: fast and efficient meta-analysis of genomewide association scans. Bioinformatics 26, 2190–1 (2010).

74. Zheng, J. et al. PhenoSpD: an integrated toolkit for phenotypic correlation estimation and multiple testing correction using GWAS summary statistics. Gigascience 7(2018).

75. Smith, S.M. et al. Enhanced Brain Imaging Genetics in UK Biobank. bioRxiv, 2020.07.27.223545 (2020).

76. Zheng, J. et al. LD Hub: a centralized database and web interface to perform LD score regression that maximizes the potential of summary level GWAS data for SNP heritability and genetic correlation analysis. Bioinformatics 33, 272–279 (2017).

77. van Rheenen, W., Peyrot, W.J., Schork, A.J., Lee, S.H. & Wray, N.R. Genetic correlations of polygenic disease traits: from theory to practice. Nature Reviews Genetics 20, 567–581 (2019).

78. Finucane, H.K. et al. Heritability enrichment of specifically expressed genes identifies disease-relevant tissues and cell types. Nat Genet 50, 621–629 (2018).

79. Tilot, A.K. et al. The Evolutionary History of Common Genetic Variants Influencing Human Cortical Surface Area. Cereb Cortex 31, 1873–1887 (2021).

80. Vermunt, M.W. et al. Epigenomic annotation of gene regulatory alterations during evolution of the primate brain. Nat Neurosci 19, 494–503 (2016).

81. Reilly, S.K. et al. Evolutionary genomics. Evolutionary changes in promoter and enhancer activity during human corticogenesis. Science 347, 1155–9 (2015).

82. Peyregne, S., Boyle, M.J., Dannemann, M. & Prufer, K. Detecting ancient positive selection in humans using extended lineage sorting. Genome Res 27, 1563–1572 (2017).

83. Vernot, B. & Akey, J.M. Resurrecting surviving Neandertal lineages from modern human genomes. Science 343, 1017–21 (2014).

84. Gazal, S. et al. Linkage disequilibrium-dependent architecture of human complex traits shows action of negative selection. Nature genetics 49, 1421–1427 (2017).

85. Gur, R.C. et al. Age group and sex differences in performance on a computerized neurocognitive battery in children age 8-21. Neuropsychology 26, 251–265 (2012).

