## Supplemental Notes and Figures for "Genome-wide association analyses of individual differences in quantitatively assessed reading- and language-related skills in up to 34,000 people"

### Supplemental Material

#### Contents

Supplemental Notes ... 3
Phenotype information ... 3
Cohort inclusion ... 3
Adolescent Brain Cognitive Development SM Study (ABCD Study) ... 4
Avon Longitudinal Study of Parents and Children (ALSPAC) ... 4
The ASTON cohort and Oxford Dyslexia cohort (ASTON and UKDYS cohorts) ... 5
Brisbane Adolescent Twin Sample (BATS) ... 5
Basque Center on Cognition, the Brain and Language cohort (BCBL) ... 5
Colorado Learning Disabilities Research Center cohort (CLDRC) ... 5
Early Language in Victoria Study (ELVS) ... 6
Familial Influences on Literacy Abilities project (FIOLA) ... 6
Generation R (GenR) ... 6
Genes, Reading, and Dyslexia study (GRaD) ... 7
Iowa Study ... 7
NeuroDys ... 7
Netherlands Twin Register (NTR) ... 7
Pediatric Imaging, Neurocognition, and Genetics cohort (PING) ... 8
The Philadelphia Neurodevelopmental Cohort (PNC) ... 8
The Raine Study ... 8
SLI consortium (SLIC) ... 8
Saguenay Youth Study (SYS) ... 9
Twins Early Development Study (TEDS) ... 9
Toronto cohort ... 9
York cohort ... 9
Evolutionary analysis ... 10
References ... 11
Supplemental Figures ... 14
Supplemental Figure 1: Manhattan plots of meta-analysis results ... 15
Supplemental Figure 2: Quantile-quantile-plots of the meta-analysis results ... 16

### Supplemental Notes

#### Phenotype information

A short survey in the GenLang network (www.genlang.org) was used to identify language- and reading-related phenotypes with prior collected data available from multiple cohorts, towards maximizing power for potential meta-analysis. The largest number of cohorts had data on word reading, nonword reading, spelling, phoneme awareness and nonword repetition. A nonword is a combination of letters that looks like a word and follows the phonotactic rules of the language, but has no meaning. In addition, performance IQ (as an index of nonverbal abilities) was analysed.

##### *Cohort inclusion*

Cohorts with language-/reading-related phenotypes and matching genome-wide genotype data were invited to join the GWAS meta-analysis effort through the GenLang network, an international consortium of researchers interested in the genomics of speech, language and reading traits. In addition, we identified a number of public cohorts with relevant phenotype and genotype data that had already been made freely available to the research community. Any cohort with measures of word reading accuracy, nonword reading accuracy, spelling accuracy, phoneme awareness and/or nonword repetition accuracy that matched the general description of these phenotypes as stated in the Methods section of the main manuscript, was invited to participate in the GWAS study. The cohorts are from Europe, North America and Australia, and include population-based, twin and disorder-oriented samples. Most cohorts were recruited in countries with English as the major native language, others are from Dutch-, Spanish-, German-, French-, Hungarian- and Finnish-speaking countries. All included phenotypes were assessed in the native language of the participants. Supplemental Table 1 provides an overview of the cohorts.

First, information from each cohort was collected on the exact phenotype measures available, including the test instrument, derived measures available, sample size, phenotype data distribution and distribution of participant age at time of data collection. A number of cohorts had measures of multiple language- and reading-related traits, sometimes measured with multiple instruments or at multiple ages. When data of multiple instruments or multiple ages were available, the instrument and/or age was used that best matched the instruments and ages collected by the other cohorts. The phenotype data distribution of all included cohorts was used to determine the phenotype data normalization method for each included trait (z-scores or rank transformation). Below you can find a description of the phenotypes included for each cohort; an overview is included in Supplemental Table 1.

Individuals over 18 years of age were excluded from all but three cohorts: the BATS cohort, PING and PNC. The PING and PNC cohorts include individuals with a wide age range: 5-21 and 8-22 years of age, respectively. The BATS cohort is a slightly older cohort, including individuals aged 11 to 26. For the BATS cohort, summary statistics were obtained as well for a subset of the cohort aged less than 18 years. Because the inclusion of the full BATS cohort did not lead to additional heterogeneity, compared to including the subset aged up to 18 years of age, the data of the full BATS cohort was used for the GWAS meta-analyses.

#### *Adolescent Brain Cognitive Development SM Study (ABCD Study)*

In the ABCD Study, word reading was assessed using the NIH Toolbox Oral Reading Recognition Test (TORRT), which measures the accuracy of pronouncing single printed words[1]. Participants can take as much time as wanted. Age-corrected standard scores were supplied. The Institutional Review Board at the University of California, San Diego approved the study, with a few sites obtaining approval of their local Institutional Review Board. All participants and/or their parents provided informed consent.

#### *Avon Longitudinal Study of Parents and Children (ALSPAC)*

ALSPAC[2,3] is a longitudinal cohort, in which reading- and language-related traits were assessed at multiple specific ages using various tools. Where possible, ages and tools were chosen that optimally matched the data of the other cohorts. The results of the Test of Word Reading Efficiency (TOWRE)[4], assessed at 13 years of age, were used as measures for word and nonword reading. The TOWRE measures the ability to sound out words and nonwords quickly and accurately, by asking participants to read through a list of words for 45 seconds, followed by a list of nonwords for 45 seconds. Spelling was assessed at 7 years of age with the Wechsler Objective Reading Dimensions (WORD) test[5], which contains 6 items assessing the ability to write letters and 44 items assessing spelling. Phoneme awareness was measured at 7 years with the Auditory Analysis Test[6], a phoneme deletion task with 2 practice and 40 test items of increasing difficulty. Participants were asked to repeat each word in full, and then with a phoneme removed. Nonword repetition was assessed using 36 nonwords of 3, 4 and 5 syllables (12 nonwords each)[7]. Participants were asked to listen to and then repeat each nonword. Performance IQ was assessed using the Wechsler Intelligence Scale for Children (WISC)[8] at 8 years of age. Please note that the study website contains details of all the data that is available through a fully searchable data dictionary and variable search tool (http://www.bristol.ac.uk/alspac/researchers/our-data/).

Pregnant women resident in Avon, UK with expected dates of delivery 1st April 1991 to 31st December 1992 were invited to take part in the study. The initial number of pregnancies enrolled is 14,541 (for these at least one questionnaire has been returned or a “Children in Focus” clinic had been attended by 19/07/99). Of these initial pregnancies, there was a total of 14,676 foetuses, resulting in 14,062 live births and 13,988 children who were alive at 1 year of age. When the oldest children were approximately 7 years of age, an attempt was made to bolster the initial sample with eligible cases who had failed to join the study originally. As a result, when considering variables collected from the age of seven onwards (and potentially abstracted from obstetric notes) there are data available for more than the 14,541 pregnancies mentioned above. The number of new pregnancies not in the initial sample (known as Phase I enrolment) that are currently represented on the built files and reflecting enrolment status at the age of 24 years is 913 (456, 262 and 195 recruited during Phases II, III and IV respectively), resulting in an additional 913 children being enrolled. The phases of enrolment are described in more detail in the cohort profile paper and its update. The total sample size for analyses using any data collected after the age of 7 years is therefore 15,454 pregnancies, resulting in 15,589 foetuses. Of these 14,901 were alive at 1 year of age.

Ethical approval for the study was obtained from the ALSPAC Ethics and Law Committee and the Local Research Ethics Committees. Consent for biological samples has been collected in accordance with the Human Tissue Act (2004). Informed consent for the use of data collected via questionnaires and clinics was obtained from participants following the recommendations of the ALSPAC Ethics and Law Committee at the time.

##### *The ASTON cohort and Oxford Dyslexia cohort (ASTON and UKDYS cohorts)*

In the ASTON and UKDYS cohorts, word reading, spelling and performance IQ was assessed using the British Ability Scale (BAS)[9]. The BAS includes an untimed single word reading test. In the spelling test, participants were asked to spell single words dictated in a sentence frame. A matrices test was used to assess performance IQ. Nonword reading was assessed using the CORE[10], lengthened to a 30-item version by Castles and Coltheart[11]. This is an untimed nonword reading task, where participants were asked to read aloud 30 nonwords consisting of one and two syllables. Phoneme awareness was assessed using a spoonerism test, where participants were asked to swap the first sounds of two words. Ethical approval was obtained from the Oxfordshire Psychiatric Research Ethics Committee and the Aston University Human Sciences Ethics Committee. All participants and/or their caregivers provided written informed consent.

##### *Brisbane Adolescent Twin Sample (BATS)*

In BATS, word and nonword reading and spelling were assessed using the CORE[10], which was lengthened to a 120-word version by Castles and Coltheart[11] to include spelling and to increase the difficulty level for an older sample. Nonword repetition was assessed using the Dollaghan and Campbell nonword repetition task[12]. Both tests were administered over the telephone by a trained researcher. The study was approved by the Human Research and Ethics Committee of the QIMR Berghofer research institute; all participants provided informed consent.

##### *Basque Center on Cognition, the Brain and Language cohort (BCBL)*

In the BCBL cohort, word and nonword reading was assessed by asking participants to read a list of 192 strings of letters that appeared in the middle of the screen, some of which were real words (96) and some of which were nonwords (96). The words included regular and irregular words. Phoneme awareness was measured using a phoneme deletion task of 24 items. Participants were asked to repeat each word without the first phoneme out loud as fast as possible. Nonword repetition was assessed by asking participants to repeat nonwords of increasing difficulty and length that were presented auditorily through headphones. A total of 24 nonwords were presented to each participant in the same order. For the reading and phoneme awareness tests, participants had a response time of 3 seconds for each item, and 5 seconds per item in the nonword repetition test. For each test, the accuracy measure was used. Performance IQ was measured with the matrices subtest of the Kaufman Brief intelligence test (K-BIT)[13]. The Institutional Review Board of the University of La Laguna approved the study and all participants provided informed consent.

##### *Colorado Learning Disabilities Research Center cohort (CLDRC)*

In the CLDRC cohort, word reading was measured with two different tests: the Time-Limited Word Recognition Test (TLWRT)[14,15] and the Word Recognition subtest of the Peabody Individual Achievement Test (PIAT)[16] word reading test. The TLWRT measures reading accuracy for words where the correct response was initiated within two seconds. The Word Recognition subtest of the PIAT is an untimed word reading test. A composite measure was used, derived in equal weights from the scores on both tests, which has proven to be a more reliable index for word reading than either measure alone[14]. Nonword reading was assessed using the TLWRT[15], similarly to the word reading assessment. Spelling was assessed using the Wide Range Achievement Test-Revisited (WRAT-R)[17], where participants were asked to write down words that were dictated. Phoneme awareness was assessed using a classical phoneme deletion task, and nonword repetition was measured using a classical nonword repetition task. Performance IQ was measured with the WISC. The study was approved by the Institutional Review Board of the University of Colorado; all participants provided informed consent.

##### *Early Language in Victoria Study (ELVS)*

ELVS is a longitudinal cohort, in which reading- and language-related traits were assessed at specific ages using various tools. Word reading and spelling was assessed using the WRAT-4[17] at 11 years of age. In the word reading subtest, participants are asked to pronounce 55 words that are presented without context. In the spelling subtest, participants are asked to write down their name, 13 letters and 42 dictated words of increasing difficulty. Participants get 10 seconds for reading each word, 5 seconds for writing letters and 15 seconds for spelling words. Nonword reading was assessed at 7 years of age using the Castle and Coltheart's reading test[11]. Phoneme awareness was measured at 5 years of age using the blending words subtest of the Comprehensive Test of Phonological Processing (CTOPP)[18], where participants have to put individual sounds together to form words. Nonword reading was measured with the Children's Test of Nonword Repetition (CNRep)[19] at 5 years of age. In this task, children are asked to repeat 40 nonwords and their accuracy is scored. Performance IQ was measured using the Wechsler Adult Intelligence Scale (WAIS)[20] at 11 years of age. Standard scores were available for all measures except nonword reading. The study was approved by the Royal Children's Hospital Human Research Ethics Committee.

##### *Familial Influences on Literacy Abilities project (FIOLA)*

In Fiola, word reading was assessed using the One-Minute-Test, or Eén-Minuut-Test in Dutch (EMT)[21], and nonword reading was assessed using the Klepel[22]. Participants were asked to correctly read as many (non)words as possible within one minute (for word reading) or two minutes (for nonword reading). The original test versions consist of a list of 116 (non)words of increasing difficulty. To avoid a ceiling effect in adults, the lists were extended by adding the last column of its parallel test, resulting in a list of 145 (pseudo)words. Phoneme awareness was measured using a phoneme-deletion test. On each test item a phoneme (always a consonant) had to be deleted from a pseudoword, resulting in another pseudoword. The test consisted of two parts. The first part comprised four monosyllabic and four disyllabic pseudowords. The second part consisted of four disyllabic pseudowords, with the phoneme that had to be deleted occurring twice. Both parts of the test started with two items (with feedback) for practice. For measuring reaction times, the timer started running after the audio file finished, until right after the participant finished pronouncing the answer. Ethical approval for this study was provided by the University of Amsterdam's Ethics Committee. Written informed consent was obtained from parents.

##### *Generation R (GenR)*

In GenR, nonword repetition was measured using the Shortened Nonword Repetition Task (NWR-S)[23], a shortened version of test developed by Rispens and Baker[24], in which 22 out of the original 40 nonwords were included. The percentage of nonwords repeated correctly was used as outcome. The study was approved by the Medical Ethical Committee of the Erasmus Medical Center in Rotterdam. All participants and/or their parents provided written informed consent.

#### *Genes, Reading, and Dyslexia study (GRaD)*

In GRaD, word and nonword reading was measured with the TOWRE[4]. Spelling was assessed with the Woodcock Johnson-III (WJ-III)[25], in which increasingly difficult words are dictated to the participant in the context of a sentence, and the participant is asked to write down the words. Phoneme awareness was measured using the CTOPP[18] elision task, during which the participants are asked to repeat a spoken word while omitting a target sound. The GRaD study was approved by the Yale Human Investigation Committee and all the review boards of participating data collection sites.

#### *Iowa Study*

In the Iowa study, word and nonword reading was measured with the Woodcock Reading Mastery Tests-Revisited (WRMT-R)[26] using the Word Identification and Word Attack subtests, respectively. In these tests, participants are asked to read aloud isolated words or nonwords from a list with increased difficulty. Spelling was assessed using the Test of Written Spelling-2 (TWS2)[27], during which participants are asked to write down dictated words with increasing difficulty. Phoneme awareness was measured using the Catts elision task[28], during which participants are asked to repeat a word while omitting a target sound. Nonword repetition was measured using the Dollaghan and Campbell nonword repetition task[12]. Performance IQ was measured with the WISC[8]. The study was approved by the Institutional Review Board of the University of Iowa IRB-01 (#200511767). All subjects in the Iowa cohort were minors who assented to participation.

#### *NeuroDys*

NeuroDys is a cohort collected in seven different European countries: Austria, Germany, Switzerland, Finland, France, Hungary and the Netherlands. For all traits, language-specific tests were used. Word reading and nonword reading was assessed by presenting language-specific lists of words or nonwords, while participants were asked to read as quickly as possible without making mistakes. Spelling was assessed by asking participants to spell single words dictated in sentence frames. Phoneme awareness was assessed using a phoneme deletion task. See [29,30] for details. Performance IQ was measured using the Block Design subtest of the WISC[8]. Ethical approval was granted by the Research Ethics Committee of the NHS (14/NS/1022), the Kantonale Ethikkomission Zürich, the Ethics Committee of the University of Salzburg, the Ethics Committee of the "Hospital District of Central Finland and the Ethics Committee of the Department of Child and Adolescent Psychiatry, Psychosomatic, and Psychotherapy of the Philipps University in Marburg. Informed written consent was given by caregivers.

#### *Netherlands Twin Register (NTR)*

The NTR is a longitudinal cohort, in which reading- and language-related traits were assessed at multiple specific ages. Where possible, ages were chosen to optimally match the data of the other cohorts. Word reading was measured using the EMT[21] and three-minutes test (Drie-Minuten Test, DMT)[31] at 9 years of age. During the DMT, participants were asked to correctly read as many words as possible within three minutes. Spelling data of children in grade 3 (comparable to grade 1 in most countries; age 6 to 7 years) was obtained from Cito's pupil monitoring system. Active spelling was measured by asking children to write down specific words dictated within sentence context. Passive spelling was measured with multiple choice questions on the spelling of a highlighted word in a sentence. Performance IQ was assessed between 6 and 18 years of age with the WISC[8], WAIS[20] or Revisie Amsterdamse Kinder Intelligentietest (RAKIT)[32]. The study was approved by the Central Ethics Committee on Research Involving Human Subjects of the VU University Medical Centre, Amsterdam, an Institutional Review Board certified by the U.S. Office of Human Research Protections (IRB number IRB00002991 under Federal-wide Assurance- FWA00017598; IRB/institute codes, NTR 03-180). Informed consent was obtained from all parents of twins.

##### *Pediatric Imaging, Neurocognition, and Genetics cohort (PING)*

In the PING cohort, word reading was assessed using the WRAT-4[17]. During this task, participants were asked to read aloud 55 words from a list. The human research protections programs and institutional review boards at the universities participating in the PING project approved all experimental and consenting procedures. Written parental informed consent was obtained for all PING subjects below the age of 18, and child assent was also obtained for all participants between the ages of 7 and 17. Written informed consent was obtained directly from all participants aged 18 years or older.

##### *The Philadelphia Neurodevelopmental Cohort (PNC)*

In the PNC, word reading was assessed using the WRAT-4[17]. Participants were asked to read aloud 55 words from a list. The institutional review boards of the University of Pennsylvania and the Children's Hospital of Philadelphia approved all study procedures. All adult participants provided informed consent; for participants under the age of 18 years, assent and parental consent were obtained.

##### *The Raine Study*

In the Raine Study, spelling was measured at 10 years of age using the Western Australian Literacy and Numeracy Assessment (WALNA). This spelling test consists of two parts: in the first part participants were asked to correct the spelling of 10 highlighted words from a written text that was also read aloud, and in the second part participants were asked to write down 14 words that were missing from written text but present when that text was read aloud. Performance IQ was assessed using Raven's Coloured Progressive Matrices (CPM)[33]. See [34] for more details. The study was approved by the Human Research Ethics Committee of King Edward Memorial Hospital, Princess Margaret Hospital, the University of Western Australia, and the Health Department of Western Australia. Consent was provided by all participants at each follow-up within the Raine Study.

##### *SLI consortium (SLIC)*

In the SLIC cohort, word reading and spelling was measured with the WORD test[5]. The word reading test consists of three parts: the first part has 4 items assessing the match between letters and beginning and end sounds, the second part has 3 items matching pictures to words, and the third part is a word reading test of 48 items increasing in difficulty. The spelling test contains 6 items to assess the ability to write letters and 44 items assessing spelling. Nonword repetition was assessed by asking participants to repeat 28 nonwords that were delivered orally or by tape. Performance IQ was assessed using the WISC[8].

Ethical permission for each collection was granted by local ethics committees. Guys Hospital Research Ethics Committee approved the collection of families from the Newcomen Centre to identify families from the South East of England with specific language disorder (Ref No. 96/7/11). Cambridge Local Research Ethics Committee approved the CLASP project "Genome Search for susceptibility loci to language disorders" (Ref No. LREC96/212). Ethical approval for the Manchester Language Study was given by the University of Manchester Committee on the Ethics of Research on Human Beings (Ref No. 03061). The Lothian Research Ethics Committee approved the project

"Genetics of specific language impairment in children in Scotland" for the use of the Edinburgh samples (Ref. No. LREC/1999/6/20). All participants provided informed consent.

##### *Saguenay Youth Study (SYS)*

In SYS, spelling was assessed using the Woodcock Johnson spelling test[25], during which participants were asked to spell and write down a list of 45 words. The number of correct answers was used as a measure for spelling. Performance IQ was measured using the WISC-III[8]. The study was approved by the Research Ethics Boards of the Chicoutimi Hospital in Chicoutimi, Quebec, Canada and the Hospital for Sick Children in Toronto, Ontario, Canada. All participants provided informed consent.

##### *Twins Early Development Study (TEDS)*

TEDS is a longitudinal cohort, in which reading- and language-related traits hav been assessed at multiple specific ages using various tools. Where possible, ages and tools were chosen that optimally matched the data of the other cohorts. Word reading and nonword reading were assessed using the TOWRE[4] at 12 years of age. Spelling was assessed using the Key stage one at 7 years of age; these data were derived from the National Pupil database. During this test, children are asked to write down 20 words that are missing from a sentence. Performance IQ was measured using the WISC[8] at 12 years of age. Ethical approval for TEDS was provided by the King's College London Ethics Committee (Ref No. PNM/09/10–104). Participants have provided informed consent at each wave of assessment.

##### *Toronto cohort*

In the Toronto cohort, word reading and nonword reading were assessed using the TOWRE[4]. Phoneme awareness was measured using the CTOPP[18] phoneme awareness composite score, a standard score based on the elision, blending words and sound matching subtests. Nonword repetition was assessed using the CTOPP nonword repetition task, during which participants are asked to repeat 18 nonwords. Performance IQ was measured using the WISC[8] III and IV. Procedural approval was given by the Hospital for Sick Children and University Health Network Research Ethics Boards. Verbal assent and/or written consent was obtained from all children and parents.

##### *York cohort*

York is a longitudinal cohort, in which reading- and language-related traits were assessed at multiple specific ages using various tools. Where possible, ages and tools were chosen that optimally matched the data of the other cohorts. Word and nonword reading were assessed using the TOWRE[4]. Spelling was assessed using the Wechsler Individual Achievement Test (WIAT)[35], during which letters, sounds and words are dictated in a sentence framework. Phoneme awareness was measured using a phoneme deletion task. Nonword repetition was assessed using the CNRep[19]. Performance IQ was measured using the WISC[8]. Ethical approval for the study was obtained from the Research Ethics Committee of the NHS (Yorkshire and the Humber – Humber Bridge) and the Ethics Committee of the Department of Psychology of the University of York. Parents provided written informed consent.

#### Evolutionary analysis

Linkage disequilibrium score regression (LDSC) heritability partitioning[36] was used to study contributions of parts of the human genome that have changed at different points during human evolution. Five different annotations were used:

- Human Gained Enhancers are regulatory regions which are active in human adult or fetal tissues and not active in macaques or chimpanzees. Thus, these regions gained regulatory function along the lineage that led to our species and hence might be involved in the emergence of human-specific traits[37,38].
- Ancient selective sweep regions consist of regions harbouring haplotypes which rose in frequency in the modern human lineage due to an advantageous allele within the haplotype in the last 300-600kya[39].
- Neanderthal-introgressed regions are the genomic variants which were introduced to the human genome by the admixture of Homo sapiens and Neanderthal populations around 50kya[40].
- Neanderthal-depleted regions are large stretches of the human genome that are depleted for Neanderthal ancestry, possibly due to critical functions and intolerance to gene flow[41].

These five evolutionary annotations were based on the annotations used previously by Tilot et al.[42] in a study of cortical surface area. We further refined those annotations by removing overlapping genomic regions among annotations. For example, we detected a small number of Neanderthal introgressed SNPs within Neanderthal lineage depleted regions, and removed these SNPs from both annotations. Further, we merged four annotations of human gained enhancers active in human foetal cortex at consecutive developmental stages. Ancient selective sweeps and adult brain-tissue expressed HGEs annotations were no different from the prior study.

### References

1. Gershon RC, Slotkin J, Manly JJ, Blitz DL, Beaumont JL, Schnipke D*, et al.* IV. NIH TOOLBOX COGNITION BATTERY (CB): MEASURING LANGUAGE (VOCABULARY COMPREHENSION AND READING DECODING). *Monographs of the Society for Research in Child Development* 2013, **78**(4)**:** 49-69.

2. Boyd A, Golding J, Macleod J, Lawlor DA, Fraser A, Henderson J*, et al.* Cohort Profile: the 'children of the 90s'--the index offspring of the Avon Longitudinal Study of Parents and Children. *Int J Epidemiol* 2013, **42**(1)**:** 111-127.

3. Fraser A, Macdonald-Wallis C, Tilling K, Boyd A, Golding J, Davey Smith G*, et al.* Cohort Profile: the Avon Longitudinal Study of Parents and Children: ALSPAC mothers cohort. *Int J Epidemiol* 2013, **42**(1)**:** 97-110.

4. Torgesen JK, Rashotte CA, Wagner RK. *TOWRE: Test of word reading efficiency*. Pro-ed Austin, TX, 1999.

5. Wechsler D. *Wechsler Objective Reading Dimensions*. The Psychological Corporation: London, 1993.

6. Rosner J, Simon DP. The auditory analysis test: An initial report. *Journal of Learning disabilities* 1971, **4**(7)**:** 384-392.

7. Gathercole SE, Willis CS, Baddeley AD, Emslie H. The children's test of nonword repetition: A test of phonological working memory. *Memory* 1994, **2**(2)**:** 103-127.

8. Wechsler D, Kodama H. *Wechsler intelligence scale for children*. Psychological corporation New York, 1949.

9. Elliott CD. *British ability scales*. nfer-nelson, 1979.

10. Bates TC, Castles A, Coltheart M, Gillespie N, Wright M, Martin NG. Behaviour genetic analyses of reading and spelling: A component processes approach. *Australian Journal of Psychology* 2004, **56**(2)**:** 115-126.

11. Castles A, Coltheart M. Varieties of developmental dyslexia. *Cognition* 1993, **47**(2)**:** 149-180.

12. Dollaghan C, Campbell TF. Nonword Repetition and Child Language Impairment. *Journal of Speech, Language, and Hearing Research* 1998, **41**(5)**:** 1136-1146.

13. Kaufman AS. *Kaufman brief intelligence test: KBIT*. AGS, American Guidance Service Circle Pines, MN, 1990.

14. Olson R, Forsberg H, Wise B, Rack J. Measurement of word recognition, orthographic, and phonological skills. 1994.

15. Olson R, Wise B, Conners F, Rack J, Fulker D. Specific deficits in component reading and language skills: Genetic and environmental influences. *Journal of learning disabilities* 1989, **22**(6)**:** 339-348.

16. Dunn LM, Markwardt FC. *Peabody individual achievement test*. American Guidance Service, Incorporated, 1970.

17. Jastak S, Wilkinson G. Wide range achievement test-revised. Wilmington, DE: Jastak Associates. *City* 1984.

18. Wagner RK, Torgesen JK, Rashotte CA, Pearson NA. *Comprehensive test of phonological processing: CTOPP*. Pro-ed Austin, TX, 1999.

19. Gathercole SE, Willis CS, Baddeley AD, Emslie H. The Children's Test of Nonword Repetition: a test of phonological working memory. *Memory* 1994, **2**(2)**:** 103-127.

20. Wechsler D. Wechsler adult intelligence scale. *Archives of Clinical Neuropsychology* 1955.

21. Brus BT, Voeten MJM. *Een-minuut-test: vorm A en B; schoolvorderingentest voor de technische leesvaardigheid, bestemd voor het tweede tot en met het zesde leerjaar van het basisonderwijs; verantwoording en handleiding*. Berkhout, 1973.

22. Van den Bos K, Spelberg HL, Scheepstra A, De Vries J. De klepel. *Vorm A en B Een test voor de leesvaardigheid van pseudowoorden Verantwoording, handleiding, diagnostiek en behandeling* 1994.

23. Clercq CMPl, Schroeff MPvd, Rispens JE, Ruytjens L, Goedegebure A, Ingen Gv*, et al.* Shortened Nonword Repetition Task (NWR-S): A Simple, Quick, and Less Expensive Outcome to Identify Children With Combined Specific Language and Reading Impairment. *Journal of Speech, Language, and Hearing Research* 2017, **60**(8)**:** 2241-2248.

24. Rispens J, Baker A. Nonword repetition: the relative contributions of phonological short-term memory and phonological representations in children with language and reading impairment. *Journal of speech, language, and hearing research : JSLHR* 2012, **55**(3)**:** 683-694.

25. Woodcock RW, McGrew, K. S., and Mather, N. *Woodcock-Johnson III*. Riverside: Itasca, IL, 2001.

26. Woodcock RW. *Woodcock reading mastery tests-revised*. American Guidance Service Circle Pines, MN, 1987.

27. Larsen SC, Hammill DD. *Test of written spelling*. Pro-ed, 1994.

28. Catts HW, Fey ME, Zhang X, Tomblin JB. Estimating the Risk of Future Reading Difficulties in Kindergarten Children. *Language, Speech, and Hearing Services in Schools* 2001, **32**(1)**:** 38-50.

29. Landerl K, Ramus F, Moll K, Lyytinen H, Leppanen PH, Lohvansuu K*, et al.* Predictors of developmental dyslexia in European orthographies with varying complexity. *J Child Psychol Psychiatry* 2013, **54**(6)**:** 686-694.

30. Moll K, Ramus F, Bartling J, Bruder J, Kunze S, Neuhoff N*, et al.* Cognitive mechanisms underlying reading and spelling development in five European orthographies. *Learning and Instruction* 2014, **29:** 65-77.

31. Verhoeven L. *Drie-minuten-toets*. Cito, 1995.

32. Bleichrodt N, Drenth P, Zaal J, Resing W. Revisie Amsterdamse Kinder Intelligentie Test. Instructie, normen, psychometrische gegevens. *Lisse: Swets en Zeitlinger* 1984.

33. Raven JC. Coloured Progressive Matrices, Sets A, A_B, B. *HK Lewis* 1962.

34. Paracchini S, Ang QW, Stanley FJ, Monaco AP, Pennell CE, Whitehouse AJ. Analysis of dyslexia candidate genes in the Raine cohort representing the general Australian population. *Genes, brain, and behavior* 2011, **10**(2)**:** 158-165.

35. Wechsler D. Wechsler individual achievement test. San Antonio, TX: Psychological Corporation; 1992.

36. Finucane HK, Bulik-Sullivan B, Gusev A, Trynka G, Reshef Y, Loh PR*, et al.* Partitioning heritability by functional annotation using genome-wide association summary statistics. *Nature genetics* 2015, **47**(11)**:** 1228-1235.

37. Vermunt MW, Tan SC, Castelijns B, Geeven G, Reinink P, de Bruijn E*, et al.* Epigenomic annotation of gene regulatory alterations during evolution of the primate brain. *Nature neuroscience* 2016, **19**(3)**:** 494-503.

38. Reilly SK, Yin J, Ayoub AE, Emera D, Leng J, Cotney J*, et al.* Evolutionary genomics. Evolutionary changes in promoter and enhancer activity during human corticogenesis. *Science* 2015, **347**(6226)**:** 1155-1159.

39. Peyregne S, Boyle MJ, Dannemann M, Prufer K. Detecting ancient positive selection in humans using extended lineage sorting. *Genome research* 2017, **27**(9)**:** 1563-1572.

40. Vernot B, Akey JM. Resurrecting surviving Neandertal lineages from modern human genomes. *Science* 2014, **343**(6174)**:** 1017-1021.

41. Vernot B, Tucci S, Kelso J, Schraiber JG, Wolf AB, Gittelman RM*, et al.* Excavating Neandertal and Denisovan DNA from the genomes of Melanesian individuals. *Science* 2016, **352**(6282)**:** 235-239.

42. Tilot AK, Khramtsova EA, Liang D, Grasby KL, Jahanshad N, Painter J*, et al.* The Evolutionary History of Common Genetic Variants Influencing Human Cortical Surface Area. *Cereb Cortex* 2021, **31**(4)**:** 1873-1887.

43. Turley P, Walters RK, Maghzian O, Okbay A, Lee JJ, Fontana MA*, et al.* Multi-trait analysis of genome-wide association summary statistics using MTAG. *Nature genetics* 2018, **50**(2)**:** 229-237.

44. Zheng J, Richardson TG, Millard LAC, Hemani G, Elsworth BL, Raistrick CA*, et al.* PhenoSpD: an integrated toolkit for phenotypic correlation estimation and multiple testing correction using GWAS summary statistics. *Gigascience* 2018, **7**(8).

### Supplemental Figures

### Supplemental Figure 1: Manhattan plots of meta-analysis results

##### Word reading

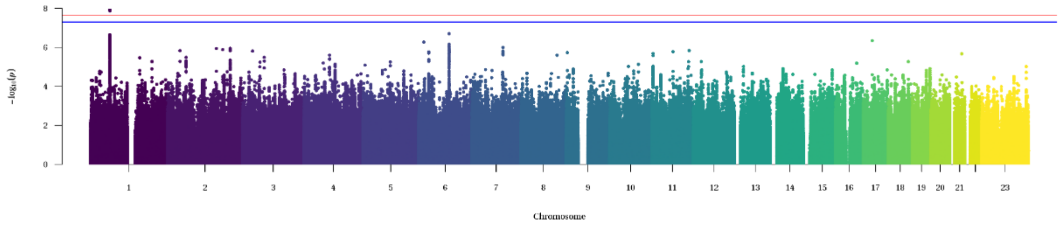

##### Spelling

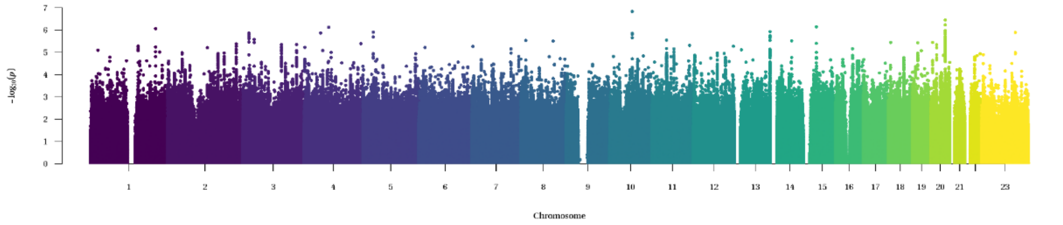

##### Phoneme awareness

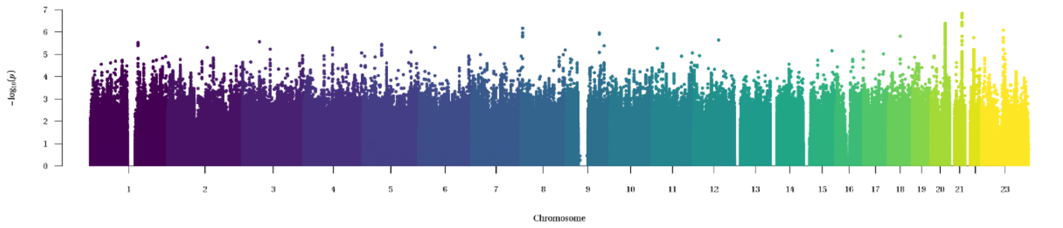

##### Non-word reading

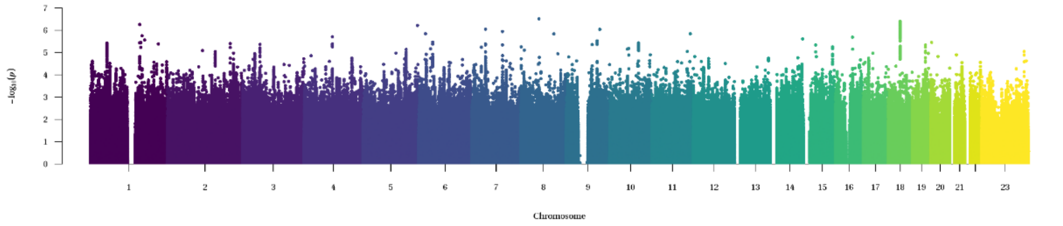

##### Non-word repetition

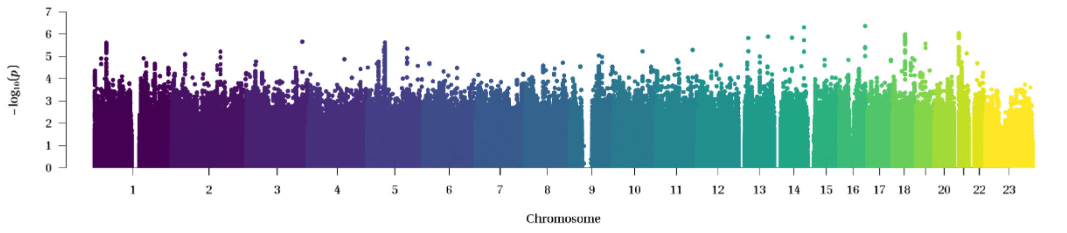

##### Performance IQ

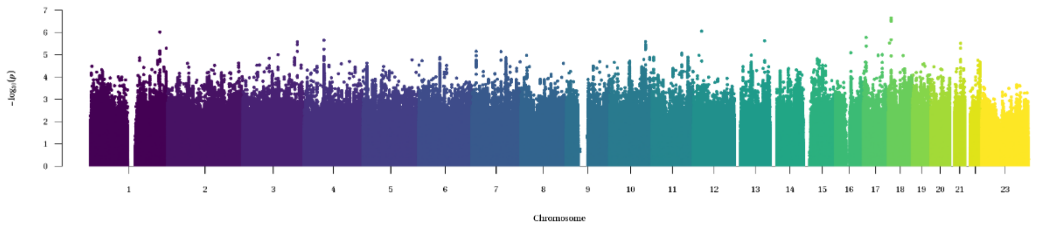

**Manhattan plots of the results of the GenLang GWAS meta-analyses.** The x axis shows chromosomal position (hg19) and the y axis represents -log(two-sided P values) for association of variants with the traits. The horizontal red line represents the threshold for genome-wide significance after correction for 2.15 independent GenLang traits ($p < 2.33 \times 10^{-8}$). The horizontal blue line represents the standard threshold for genome wide significance ($p < 5 \times 10^{-8}$).

#### Supplemental Figure 2: Quantile-quantile-plots of the meta-analysis results

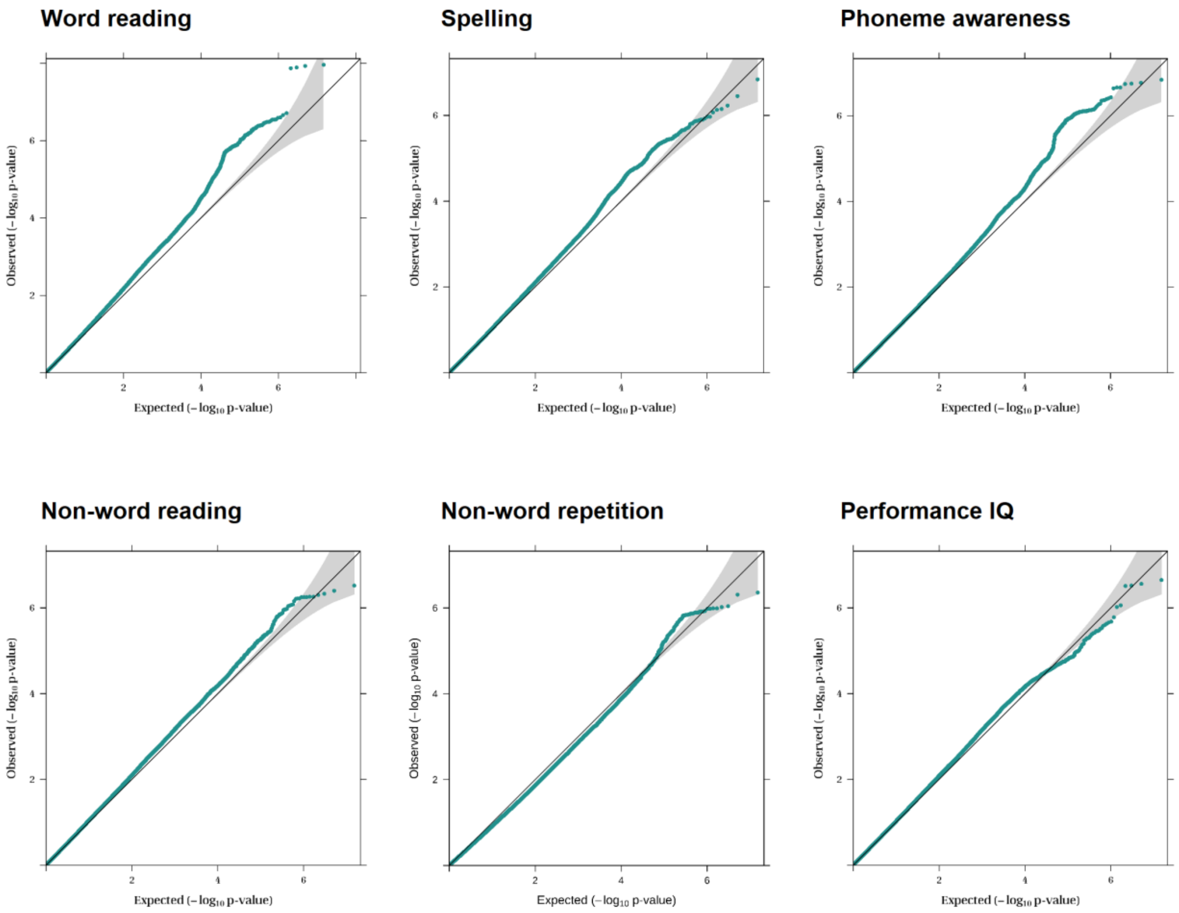

**Quantile-quantile plots of each of the GenLang meta-analyses.** The gray-shaded areas in the plots represent the 95% confidence intervals under the null hypothesis.

Supplemental Figure 3: Quantile-quantile-plots of the heterogeneity analyses of the meta-analyses.

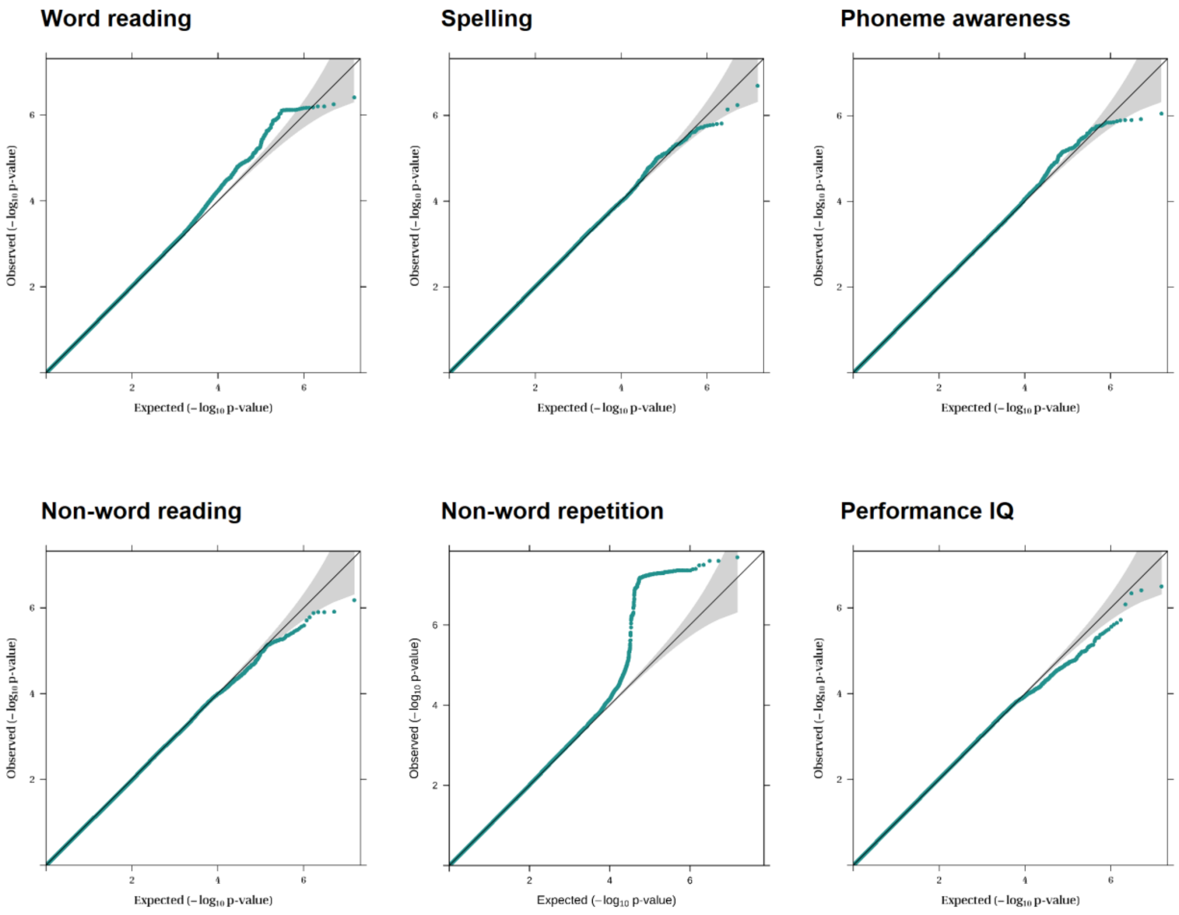

**Quantile-quantile plots of the Cochran's Q-test p-values for each of the GenLang meta-analyses.** The gray-shaded areas in the plots represent the 95% confidence intervals under the null hypothesis. For nonword repetition, a random-effects meta-analysis was performed because of the high heterogeneity.

### Supplemental Figure 4: Manhattan plots and quantile-quantile plots of the multivariate word reading results

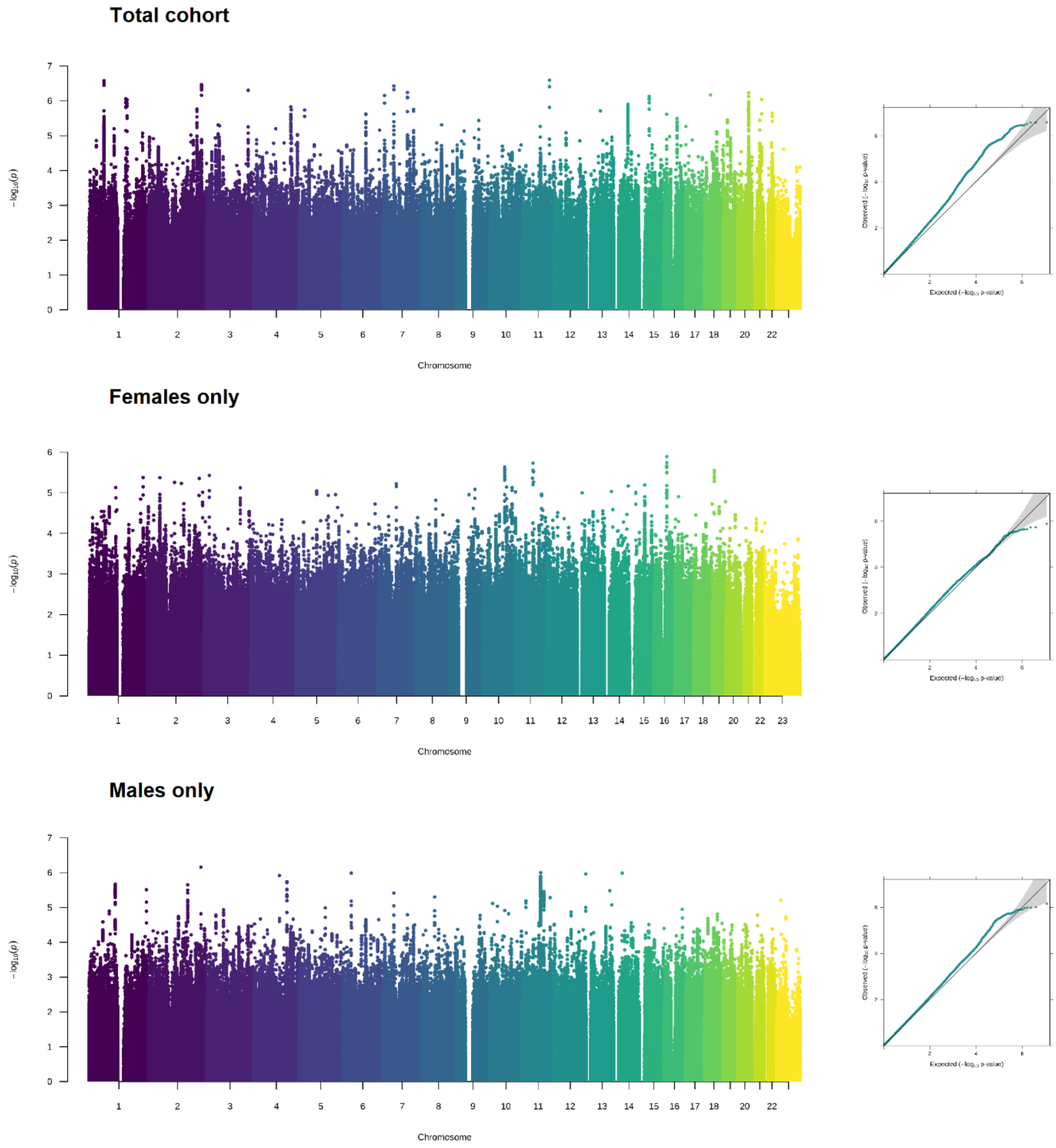

**Manhattan and quantile-quantile plots of the multivariate word reading results.** The multivariate GWAS analysis was performed using MTAG[43] to improve the power of the univariate word reading GWAS meta-analysis by incorporating information of three highly correlated traits: non-word reading, spelling and phoneme awareness. Left: Manhattan plot of the p-values of the MTAG multivariate GWAS analysis. The y axis represents -log(two-sided P values) for association of variants with the traits. Right: Quantile-quantile plots of the p-values of the MTAG multivariate GWAS analysis. The gray-shaded areas represent the 95% confidence intervals under the null hypothesis. The plots are shown of the MTAG results for the total cohort and for female-only and male-only analyses.

### Supplemental Figure 5: Locuszoom plot and forestplot of the chromosome-1 locus significantly associated with word reading

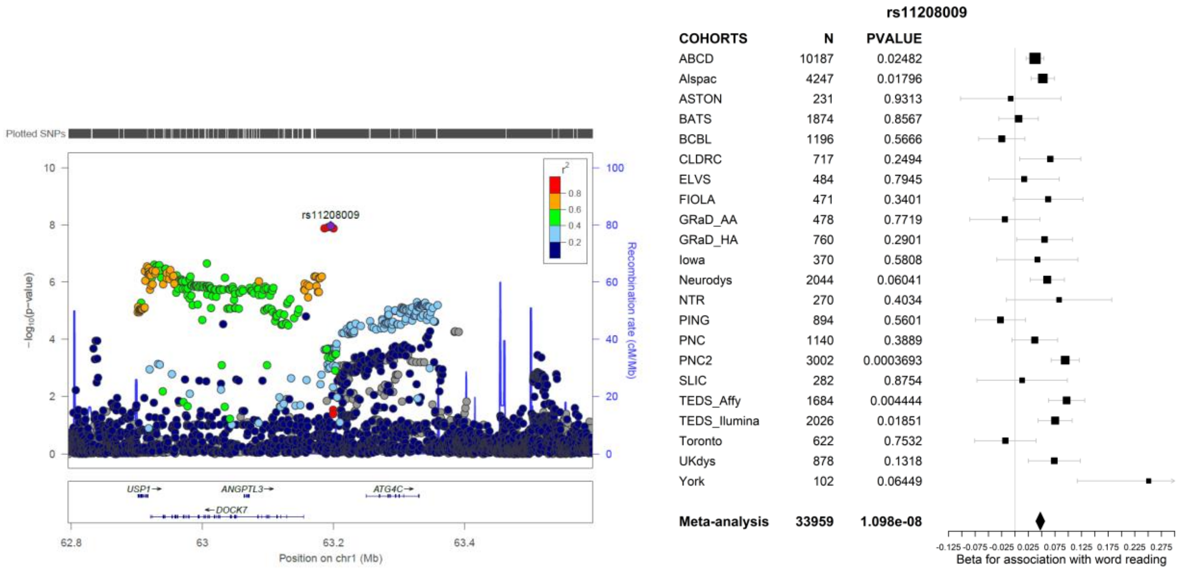

**Genome-wide significant locus associated with word reading.** Left: Locuszoom plot. Colours represent linkage disequilibrium with rs11208009 based on the 1000 Genomes project reference data. Right: Forest plot of the association results for rs11208009 in each of the GenLang cohorts.

Supplemental Figure 6: Overview of surface-based morphometric (SBM) neuroanatomical traits, assessed via brain imaging, and their genetic correlations with the multivariate word reading results.

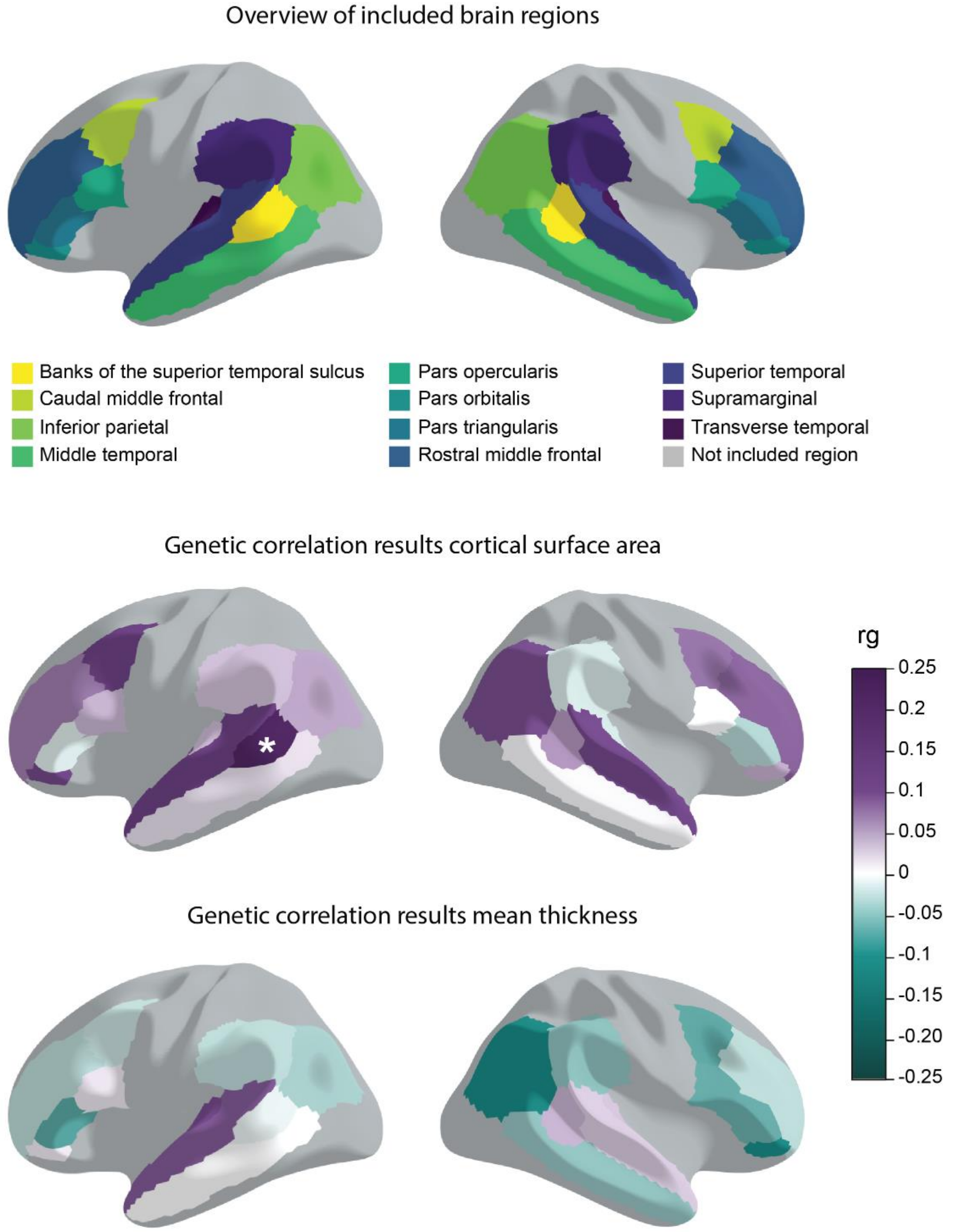

Top: Overview of the cortical areas included in the genetic correlation analysis with the multivariate word reading results. The 11 regions were selected based on a literature review encompassing brain regions and white matter tracts with known links to aspects of reading and language (see Methods). For each of the 11 regions, summary statistics of cortical surface area and mean thickness from the UK Biobank were obtained for the left and right hemisphere. Genetic correlations (rg) with the multivariate word reading results were estimated with LD score regression. Results are shown for cortical surface (middle) and mean thickness (bottom). Purple to green colours represent genetic correlations (rg) with the multivariate word reading results. Grey areas are not included in the genetic correlation analyses. *P < $1.89 \times 10^{-3}$ ($p<0.05$ after correction for 26.49 independent brain imaging traits). The cortical surface area of the banks of the superior temporal sulcus of the left hemisphere is significantly correlated with the multivariate word reading results. Results can also be found in Supplemental Table 14.

#### Supplemental Figure 7: Overview of diffusion tensor imaging (DTI) traits and their genetic correlation with the multivariate word reading results.

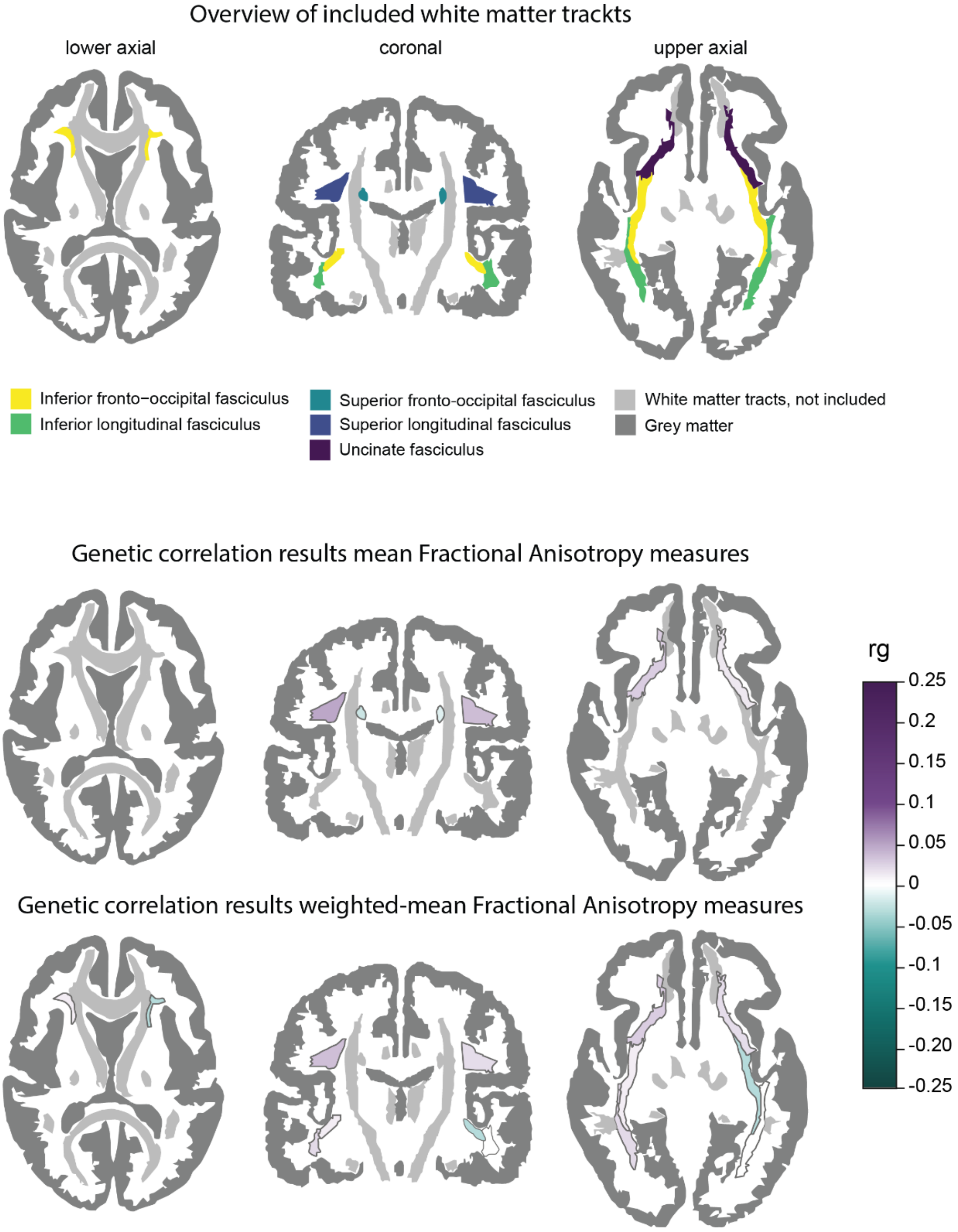

Top: Overview of the DTI traits included in the genetic correlation analysis with the multivariate word reading results. Included traits are summary statistics of 14 DTI traits of five different white matter tracts from the UK Biobank, selected based on a literature review encompassing brain regions and white matter tracts with known links to aspects of reading and language (see Methods). Middle and bottom: overview of the genetic correlations (rg) between the multivariate word reading GWAS meta-analysis results and the DTI traits, estimated with LD score regression. Separate plots are made for the mean Fractional Anisotropy (FA) measures (middle) and the weighted-mean FA measures (bottom). Results can also be found in Supplemental Table 14.

#### Supplemental Figure 8: Genetic correlations between UK Biobank brain imaging traits analysed in this study.

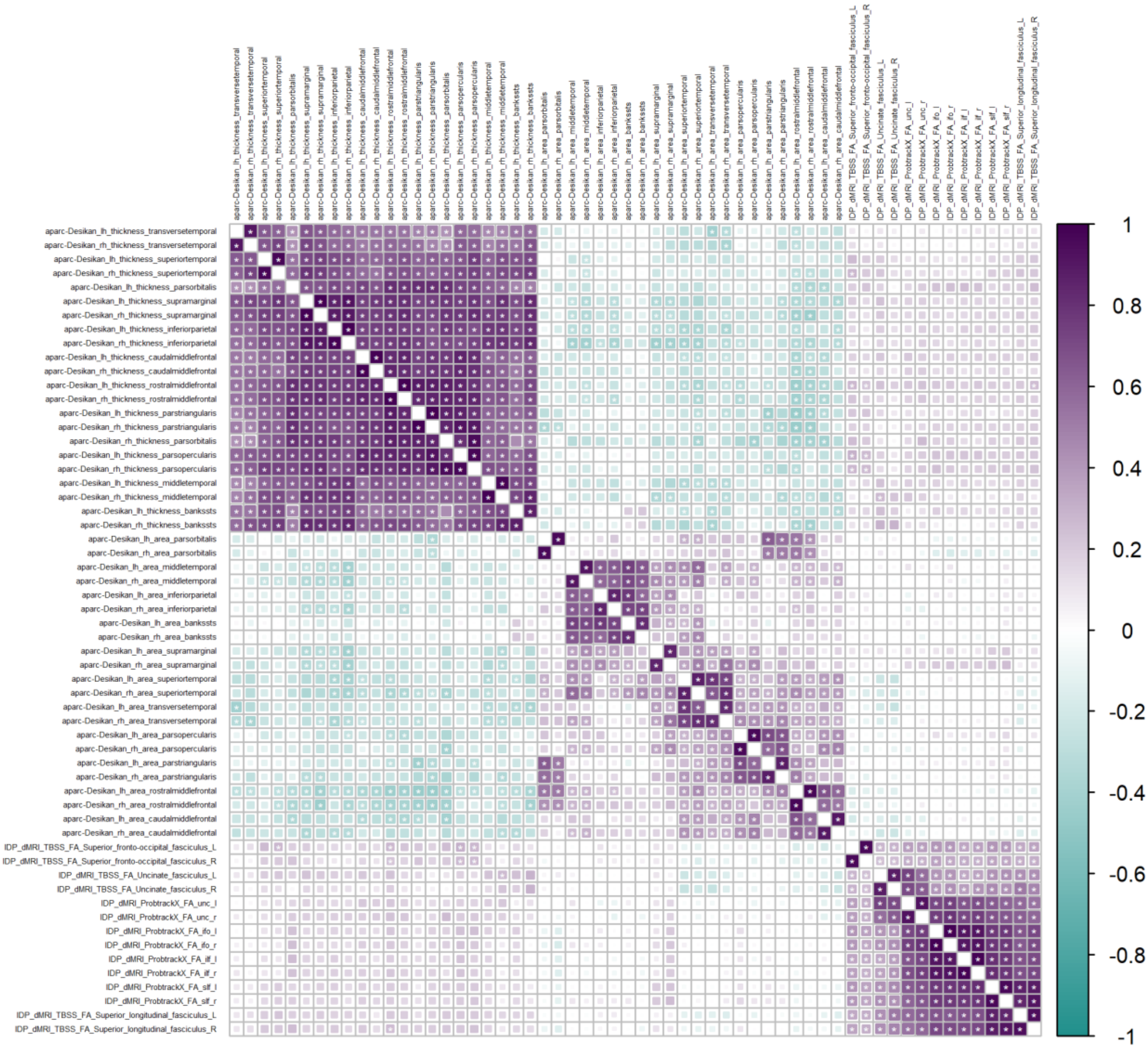

Genetic correlations (rg) between the 58 UK Biobank derived brain imaging traits were calculated with LDSC. **Significant correlation after correction for 58*57/2 tests ($p < 3.02 \times 10^{-5}$). The correlation matrix was used as input for PhenoSpD[44] to calculate the effective number of independent variables.

#### Supplemental Figure 9: LDSC heritability partitioning identifies significant enrichment in three annotations reflecting brain enhancers

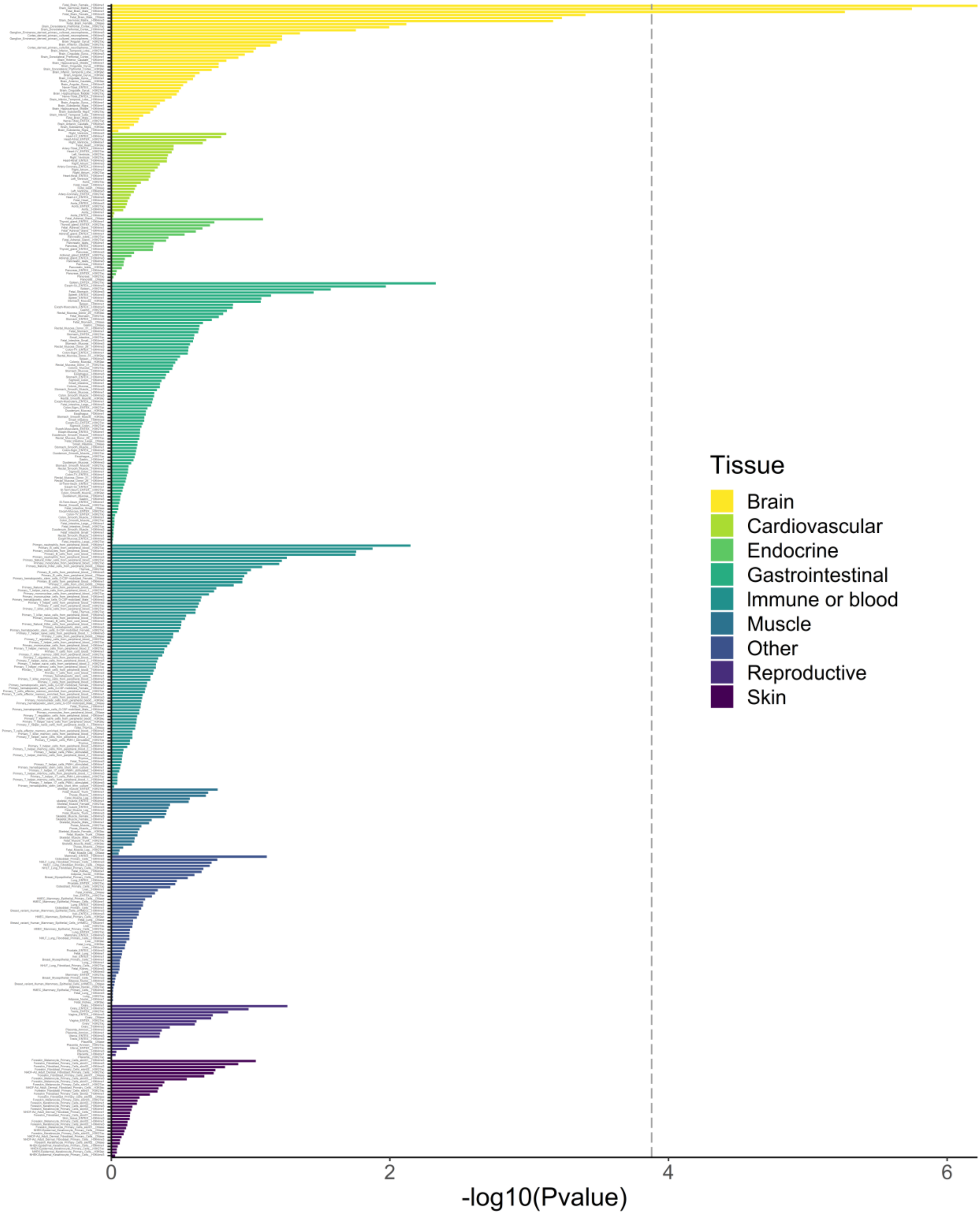

**LDSC heritability partitioning of MTAG results for 489 annotations of tissue-specific chromatin signatures.** The graph shows -log-10 p-values on the y-axis and brain region-specific chromatin mark on x-axis. The dashed line shows the p-value threshold for significant enrichment after Bonferroni-correction for testing 489 annotations. The results are available as a table in Supplemental Table 17.

Supplemental Figure 10: Gene property analysis with MAGMA identifies no significant relation between the MTAG results and tissue-specific expression

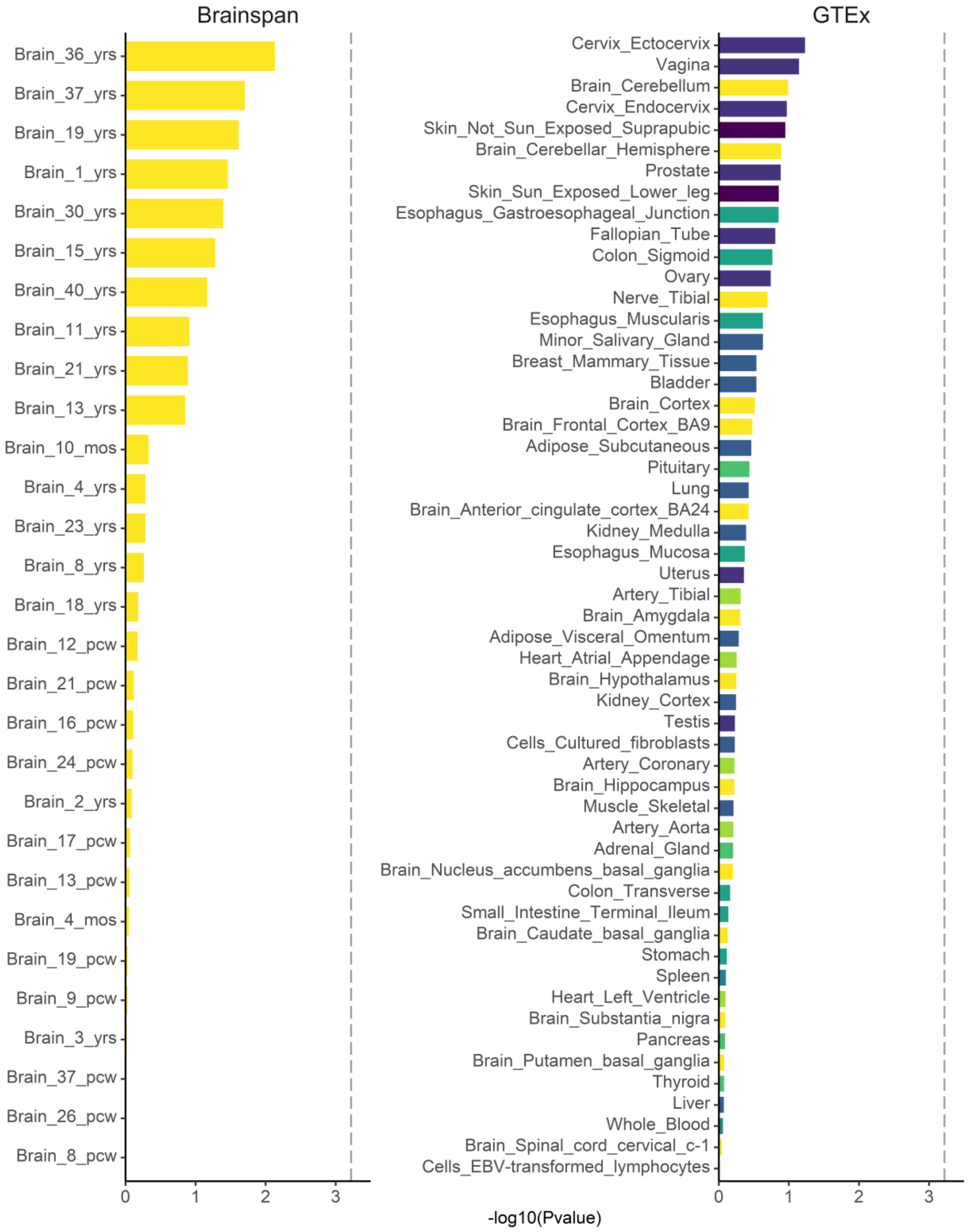

**MAGMA gene property analysis of the multivariate word reading results using two RNA-sequencing datasets of human tissue**. Brainspan data includes human brain tissue at different ages during development, and GTEx V8 data includes a wide range of adult human tissues. The graph shows -log-10 p-values on the y-axis and cell types on the x-axis. The dashed line shows the p-value threshold for significant enrichment after Bonferroni correction for testing 83 tissues.

Supplemental Figure 11: Gene property analysis with MAGMA identifies a relation between the MTAG results and embryonic red nucleus neurons

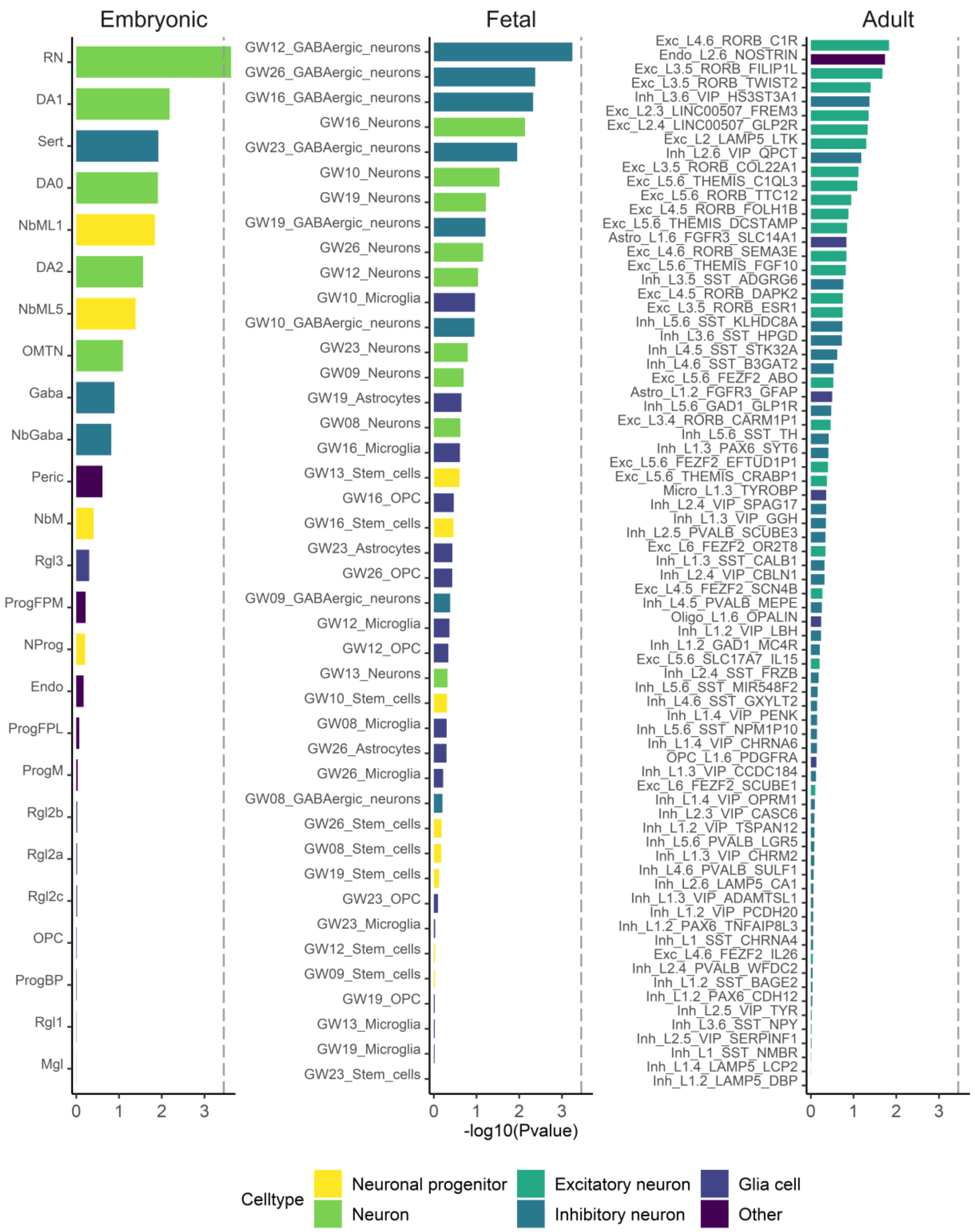

**MAGMA gene property analysis using three single-cell RNA-sequencing datasets of human brain tissue, representing embryonic midbrain and fetal and adult cortex.** The graph shows -log-10 p-values on the y-axis and cell types on the x-axis. The dashed line shows the p-value threshold for significant enrichment after Bonferroni correction for testing 142 cell types.
